# Predicting and validating protein degradation in proteomes using deep learning

**DOI:** 10.1101/2020.11.29.402446

**Authors:** Matiss Ozols, Alexander Eckersley, Christopher I. Platt, Callum S. McGuinness, Sarah A. Hibbert, Jerico Revote, Fuyi Li, Christopher E.M. Griffiths, Rachel E.B. Watson, Jiangning Song, Mike Bell, Michael J. Sherratt

## Abstract

Age, disease, and exposure to environmental factors can induce tissue remodelling and alterations in protein structure and abundance. In the case of human skin, ultraviolet radiation (UVR)-induced photo-ageing has a profound effect on dermal extracellular matrix (ECM) proteins. We have previously shown that ECM proteins rich in UV-chromophore amino acids are differentially susceptible to UVR. However, this UVR-mediated mechanism alone does not explain the loss of UV-chromophore-poor assemblies such as collagen. Here, we aim to develop novel bioinformatics tools to predict the relative susceptibility of human skin proteins to not only UVR and photodynamically produced ROS but also to endogenous proteases. We test the validity of these protease cleavage site predictions against experimental datasets (both previously published and our own, derived by exposure of either purified ECM proteins or a complex cell-derived proteome, to matrix metalloproteinase [MMP]-9). Our deep Bidirectional Recurrent Neural Network (BRNN) models for cleavage site prediction in nine MMPs, four cathepsins, elastase-2, and granzyme-B perform better than existing models when validated against both simple and complex protein mixtures. We have combined our new BRNN protease cleavage prediction models with predictions of relative UVR/ROS susceptibility (based on amino acid composition) into the Manchester Proteome Susceptibility Calculator (MPSC) webapp http://www.manchesterproteome.manchester.ac.uk/#/MPSC (or http://130.88.96.141/#/MPSC). Application of the MPSC to the dermal proteome suggests that fibrillar collagens and elastic fibres will be preferentially degraded by proteases alone and by UVR/ROS and protease in combination, respectively. We also identify novel targets of oxidative damage and protease activity including dermatopontin (DPT), fibulins (EFEMP-1,-2, FBLN-1,-2,-5), defensins (DEFB1, DEFA3, DEFA1B, DEFB4B), proteases and protease inhibitors themselves (CTSA, CTSB, CTSZ, CTSD, TIMPs-1,-2,-3, SPINK6, CST6, PI3, SERPINF1, SERPINA-1,-3,-12). The MPSC webapp has the potential to identify novel protein biomarkers of tissue damage and to aid the characterisation of protease degradomics leading to improved identification of novel therapeutic targets.

## Introduction

Although the causative mechanisms of ageing are not yet fully understood, there is compelling evidence that biochemical pathways, including protein oxidation, proteolysis and protease-mediated cleavage, contribute to loss of proteostasis (protein homeostasis) and the age-related decline of organs (1). Similarly, oxidative stress and aberrant protease activity are also implicated in pathological remodelling of both acute and chronically inflamed tissues and organs (2–4).

In skin, ultraviolet radiation (UVR), oxidative stress and upregulated protease activity are interlinked processes associated with clinical photoageing which manifests in the dermis as profound histological remodelling of fibrillar collagen and elastic fibres and the accumulation of protein carbonyls, oxidative damage (5–8). Collectively this UV, ROS and protease driven proteolysis in ageing and diseased tissue can be termed the degradome (9). A better understanding of the degradomic degeneration of tissue proteomes may lead to identification of novel biomarkers of damage and bioactive matrikines which can be used to design novel therapeutics (10,11).

Previous studies from our group suggest that both the *in vivo* remodelling of the elastic fibre-associated fibrillin microfibrils, which is a hallmark of early photoageing, and the relative molecular and ultrastructural susceptibility of these assemblies *in vitro* to UVR is likely to be due to specific amino acid (AA) compositions of major components such as fibrillin-1 (7,12,13). These proteins, and others also enriched in both UVR-absorbing (UV-chromophore) and oxidation-sensitive AA residues (e.g. fibronectin and lens crystallins), are susceptible to degradation by environmentally attainable UVR doses (14). In contrast, these doses and wavelengths do not affect the electrophoretic mobility of collagen I or tropoelastin, which are largely devoid of these AA residues, or the ultrastructure of collagen VI which contains fewer UV-chromophores than either fibronectin or fibrillin-1 (13). Recently we have also demonstrated, using a newly-developed proteomic peptide location fingerprinting methodology, that UVR exposure can induce subtle structure-associated changes in the largest collagen VI alpha chain (alpha-3) which are not detectable as changes in global collagen VI microfibril ultrastructure (15) or architecture (16). Therefore, AA composition appears to be a good predictor of relative susceptibility to UVR and oxidative damage (which is also a factor in non-UVR exposed tissues) (17–20). However, predicting the relative susceptibility of proteins to protease-mediated cleavage is a more difficult task, as enzymatic proteolysis is dependent on not only the primary structure (i.e. AA sequence) but also on protein folding and interactions between enzymes and the exposed protein surface, and hence 2-dimensional (D), 3D or quaternary structure (21).

The matrix metalloproteinases (MMPs) are a large family of zinc-dependent enzymes which are thought to play key roles in skin photoageing and many other age- and disease-related disorders (22). Other enzyme families such as serine proteases can also degrade ECM proteins (23,24) but their action in skin ageing is not as well characterised. For enzymes such as trypsin (where cleavage occurs at the C-terminal side of Arg and Lys AA residues when not followed by Pro) and the *Staphylococcus aureus* protease V8 - GluC (where cleavage occurs on the C-terminal side of Glu in preference to Asp), the validated prediction of cleavage sites is well established (25,26). However, the prediction of cleavage sites in ECM proteins by proteases such as the MMPs and cathepsins requires the development and application of complex mathematical models based on state-of-the-art machine learning and deep learning techniques which utilise data available in databases such as MEROPS (27,28). A number of bioinformatic tools have been developed to predict cleavage sites in AA sequences. For example, PROSPER, which is based on Support Vector Machines (SVMs), utilises some structural features to perform the prediction and DeepCleave, developed more recently, uses state-of-the-art deep convolutional neural network (CNN) (29–32). However, to our knowledge the predictions of these algorithms have not been experimentally validated against native ECM proteins, nor has their comparative performance been evaluated in this context. Moreover, although recurrent neural networks have shown promising results in protein sequence and function prediction (33), this architecture has not previously been evaluated for proteolytic cleavage site prediction (34,35), to our knowledge.

This first goal of this study was to use the AA sequence information and relevant sequence-derived features to: i) predict and stratify the relative susceptibility of dermal ECM proteins (as defined by the Manchester Skin Proteome (36)) to UVR/oxidative damage; ii) relate these predictions to published experimental data; and iii) identify new potential targets of photodamage and photo-oxidation. The second goal was to test the ability of PROSPER and DeepCleave to identify MMP9-determined cleavage sites in two exemplar purified proteins (decorin [DCN] and vitronectin [VTN]) and subsequently in a complex ECM proteome derived from cultured human dermal fibroblasts (HDFs). Given the relatively poor performance of these algorithms against native ECM proteins, our third aim was to develop and evaluate a new protease cleavage prediction algorithm: Manchester Proteome Cleave (MPC) which uses state-of-the-art deep bidirectional recurrent neural network (deep BRNN) architecture (34,35). Finally, our last aim was to integrate both UVR/ROS and protease prediction methods into a webtool, termed the ‘Manchester Proteome Susceptibility Calculator (MPSC)’ which can predict the relative susceptibility of proteins to degradative mechanisms and hence identify potential novel biomarkers of age- and disease-induced tissue damage.

## Results

### Predicted UVR/ROS susceptibilities correlate well with published experimental data and reveal novel potential markers of photodegradation

Our first aim was to survey the amino acid composition of all skin proteins to predict relative UVR/ROS susceptibility. Ultraviolet radiation can be subdivided into three distinct wavelength ranges - UVC (100–280 nm), UVB (280–315 nm), and UVA (315–400 nm); UVC is absorbed by the ozone layer, and so solar UVR at the Earth’s surface consists of 5% UVB and 95% UVA. These wavelengths can penetrate skin, generating reactive oxygen species (ROS) (7,37). The AA residues Trp, Tyr and double-bonded Cys (Cys=Cys) are sensitive to both biologically relevant wavelengths of UVR and oxidation (14), whereas Met, His, and Cys are sensitive to oxidation alone (38). We have previously established that proteins rich in these AA residues are at risk of degradation by UVR and/or ROS (7). In this study, building upon our initial insights, we have established an effective mathematical model allowing the prediction of relative protein susceptibilities to UVR/ROS within a whole skin proteome. To validate the UV/ROS model, we reviewed published data from experimental studies which subjected ECM proteins and assemblies to physiologically relevant UVR doses and wavelengths and categorised selected proteins as susceptible, semi-susceptible or resistant accordingly **(Supporting File 1)**. We show that the mathematical model correctly predicts the relative experimental susceptibilities of these proteins **(Fig.1.a)**. We have integrated this mathematical model in our webtool, the MPSC, which allows analysis of protein sequences for their UV, ROS and UV/ROS susceptibilities.

**Figure 1.**
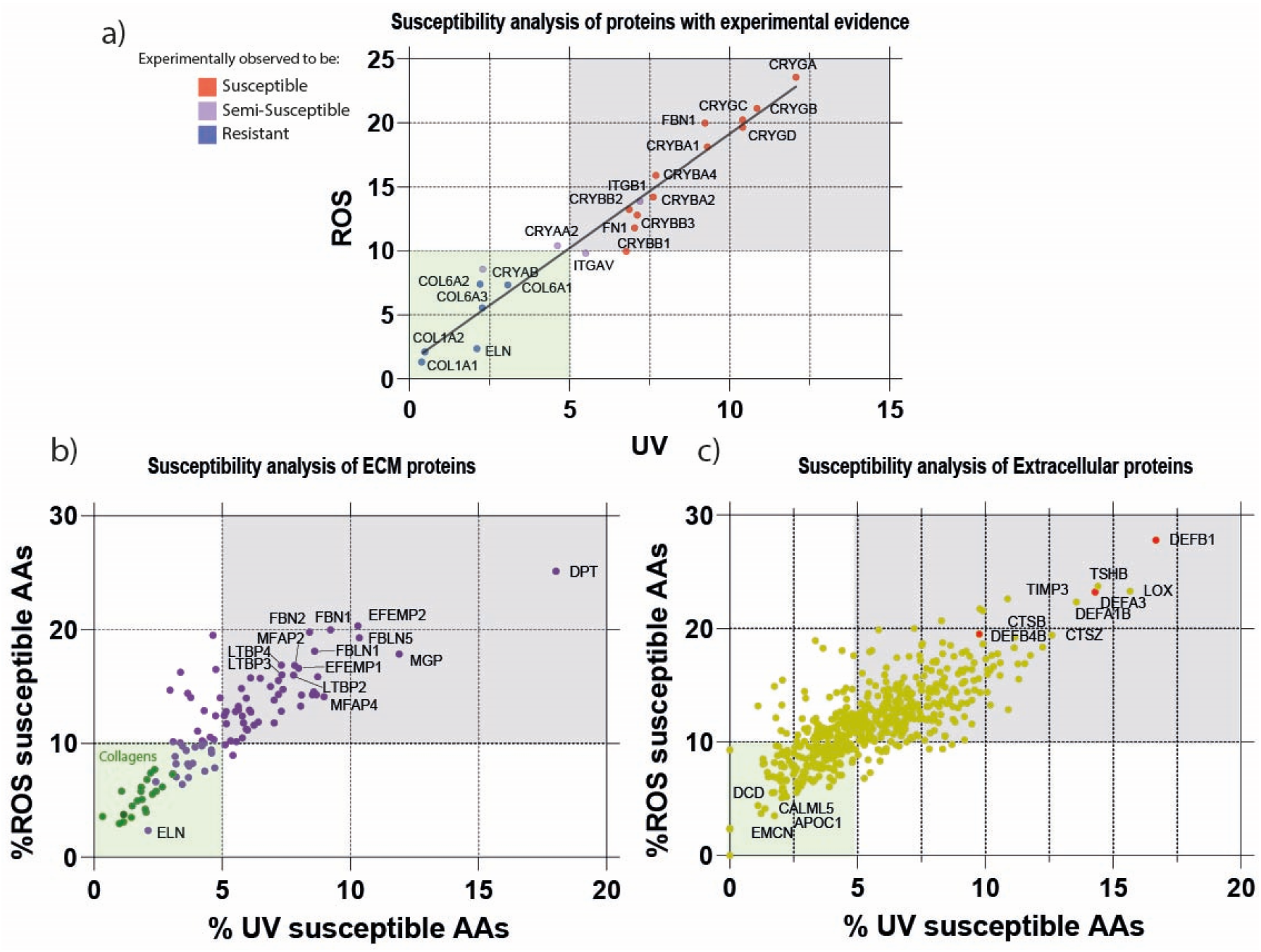
Predicted susceptibilites of skin proteins to photo- and oxidative-damage based on AA composition. a) Predicted UVR and/or ROS susceptibilities of proteins previously reported as susceptible or resistant by experimental observations. Experimentally verified UVR and ROS susceptible proteins (red) are enriched in UVR- and ROS-susceptible moieties compared with resistant proteins (blue). The predicted UVR- and ROS-susceptibilities exhibited a positive linear correlation (R^2^ = 0.94). Applying thresholds to distinguish between experimentally susceptible and resistant proteins suggest that a composition of > 5% UVR AA residues and > 10% oxidation-sensitive AA residues may be indicative of UV/ROS susceptibility (susceptible = grey box, resistant = green box). b) Predicted UVR- and ROS-susceptibilities of skin’s ECM proteins. Elastic fibre-associated proteins, except for elastin itself, and collagens (green) clearly stratify into susceptible and resistant risk categories, respectively. c) Predicted UVR and ROS susceptibility of non-ECM extracellular proteins. Defensins (red) are highlighted as an example of a susceptible protein family.

Applying the MPSC to ECM proteins in the (as defined by the Manchester Skin Proteome) we predict that many elastic fibre-associated proteins which play roles in fibre formation, structure and organisation (FBN1, FBN2, FBLN1, FBLN5, LTBP2, LTBP3, LTBP4, EFEMP2, MFAP2 and MFAP4) (39,40), will be highly susceptible to degradation by both UVR and oxidation **(Fig.1 b)**. These predictions agree well with *in vivo* observations of dermal remodelling in photoageing. For example, a key hallmark of severely photoaged skin is the presence of disorganised material containing multiple elastic fibre proteins, termed solar elastosis (41) whilst mildly photoaged skin is characterised by the loss of FBN1 and FBLN5 microfibrils from the upper dermis which is exposed to the highest UVR dose(42,43). We have demonstrated *in vitro* that chromophore-rich fibrillin-1 (in the form of fibrillin microfibrils) is susceptible to low dose UVR whilst chromophore-poor tropoelastin (ELN; the elastin precursor) is resistant (12,13). Our analysis predicts that skin collagens would be relatively resistant to both UVR and to ROS-mediated oxidation **(Fig.1a&b)**. Previously we have shown that the eletrophoretic mobility of collagen I, the most abundant skin protein, is unaffected by physiolgically relevant doses of UVR (12). In contrast, it is clear from observational studies that collagen I abundance decreases in aged (and particularly in photo-aged) human skin, which leads to dermal thinning (44,45). In additon to predicting the relative UVR/ROS suceptibilties of major structural components such as elastin and the collagens, our analysis also identifies potential novel markers involved in the regulation of TGFβ and collagen fibril formation (DPT) and ECM development (MGP) which, to our knowledge, have not previously been implicated in photoageing **(Fig.1.b)**.

Extracellularly, MPSC predicts that the defensin family (DEFB1, DEFA3, DEFA1B and DEFB4B) of proteins may be UVR/ROS susceptible **(Fig.1c)**. This is consistent with the ability of defensins to act as antioxidant proteins (46). Crucially, MPSC not only predicts that known photoageing biomarkers (LOX and TIMP3) will be susceptible to UVR/ROS degradation (47–49) but also highlights the potential susceptibility (TSHB, CTSB, CTSZ) and resistance (DCD, EMCN, CALM5 and APOC1) of new skin ageing biomarker proteins **(Fig.1.c)**.

Whilst UVR/oxidative damage is clearly an important mediator of protein degradation in skin, protease-induced proteolysis is also thought to play a key role. Degradative mechanisms do not act in isolation (50,51) and we, and others, have shown that these two mechanisms may interact with UVR exposure to enhance subsequent protease action (7,52). Our prediction that dermal collagens I, III, IV, V and VII are likely to be highly resistant to UVR and ROS implies that their degradation *in vivo* must be mediated by other mechanisms such as extracellular proteases.

### Accurate prediction of protease cleavage sites in native ECM proteins is challenging

We next aimed to predict the location and number of protease cleavage sites within skin-expressed proteins. As the relative abundance of AA residues can predict protein susceptibility to UVR/ROS, we hypothesise that protease cleavage site load can determine the relative susceptibility of proteins to enzymatic degradation. In skin, MMPs and other proteases such as elastase and members of the cathepsin and granzyme families play a significant role in tissue maintenance and remodelling (50,51). Prediction of proteolytic cleavage sites is dependent on not only the 1° AA sequence but also the higher order 2°-4° structures which determine features such as solvent accessibility and disordered regions (53,54). Although currently available algorithms such as PROSPER and DeepCleave can predict cleavage sites with a good degree of accuracy (29,30), these predictions are: i) often limited to a specific subset of proteases, and; ii) lack experimental validation both in a simple and complex model systems. In this study, we used SDS-PAGE and mass spectrometry to characterise protein degradation and to validate and compare protease cleavage site predictions (by PROSPER and DeepCleave) for two purified native ECM proteins.

Initially, we used a simple model system, digesting purified DCN and VTN with recombinant MMP9 (chosen as an exemplary enzyme due to its importance in skin photoageing process (55)). By gel-electrophoresis we confirmed that both DCN and VTN are MMP9 substrates (56,57). In quadruplicate experiments, following MMP9 exposure, the band corresponding to VTN was absent and there was evidence of substantial aggregation. In contrast, the DCN band remained detectable following MMP9 exposure although there was some evidence of degradation **(Fig.2.a) (Supporting File 2)**. Cleavage sites in these two proteins were characterised experimentally by performing subsequent in-gel digestion of the same gels with trypsin and liquid chromatography tandem mass spectrometry (LC-MS/MS) followed by a bioinformatic analysis pipeline which searched for non-tryptic (assumed to be MMP9-derived) sites in the digested samples. Using LC-MS/MS, we identified 33 putative cleavage sites in VTN compared with 18 in DCN **(Fig, 2b)**. LC-MS/MS revealed that cleavage sites in VTN were predominantly densely packed between AAs 300-400, corresponding to the heparin-binding domain which has proven involvement in fibronectin deposition (58). In contrast, DCN cleavage sites were distributed throughout the AA sequence **(Fig.2.b)**. We next compared these experimentally determined cleavage sites with those predicted by PROSPER and DeepCleave. While PROSPER and DeepCleave achieve excellent performance against data available in MEROPS, their accuracy in predicting MMP9 cleavage sites in native VTN and DCN (as evaluated by AUC score) was only slightly better than random (PROSPER AUC: 0.55; DeepCleave AUC: 0.64) **(Fig.2.c)**.

**Figure 2.**
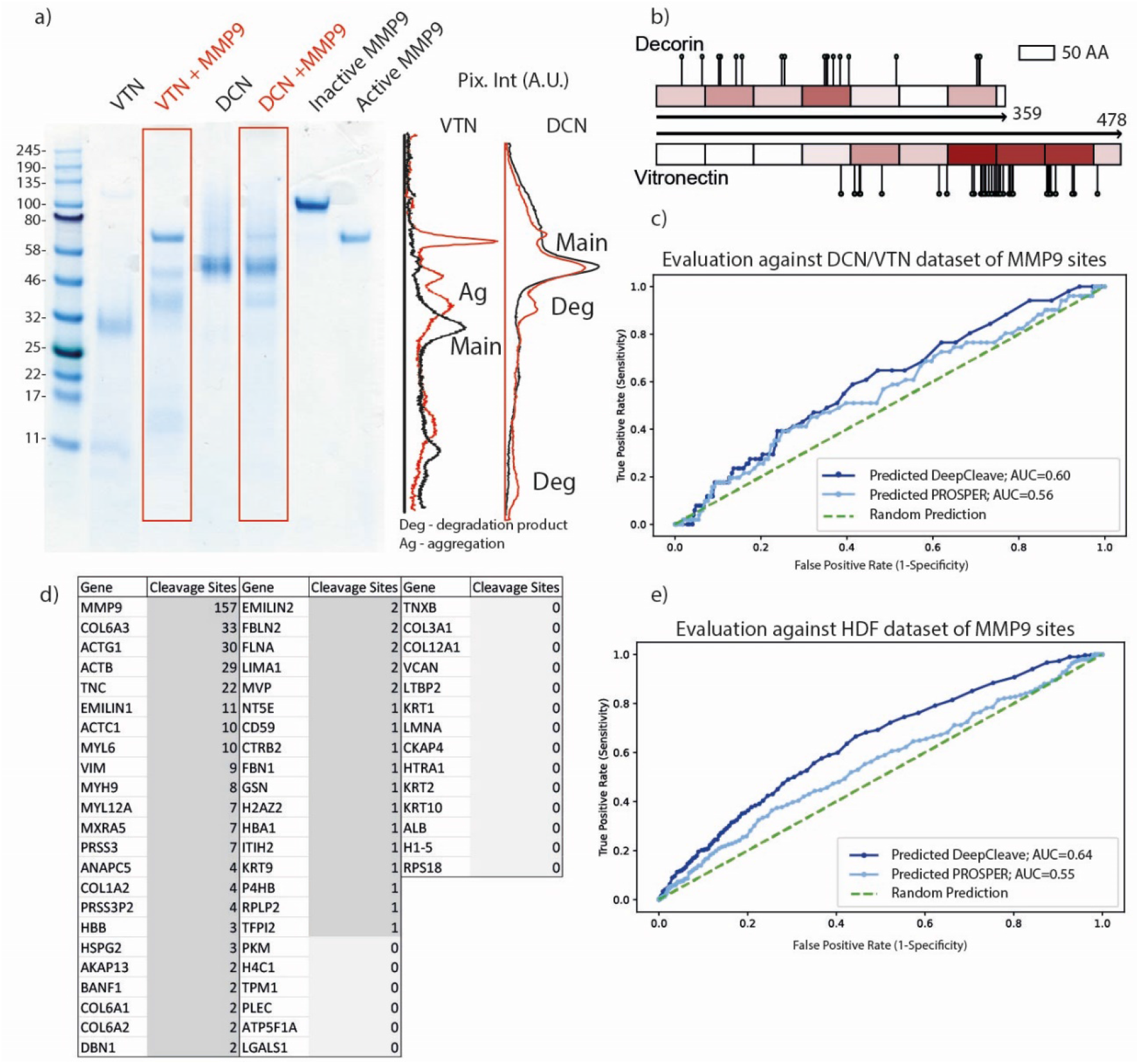
Prediction and validation of protease cleavage sites. a) Gel-electrophoresis and associated densitometry profiles for control (black) and MMP9-digested (red) DCN and VTN proteins (although the predicted molecular weight (mw) of this recombinant DCN is 39 kDa, as a the attached glycosaminoglycan chain shifts it mw to 50 kda, as described by the manufacturers). MMP9 exposure led to the disappearance of the VTN main band (~30 kDa) and the formation of a heavier protein aggregation band (~45 kDa) whereas MMP9 exposure led to a reduced staining of the DCN main band density and the formation of a lower molecular weight degradation product. This suggests that VTN may be more susceptible to MMP9 than DCN. b) Differential degradation of VTN and DCN was confirmed by LC-MS/MS on bands excised from same gel. Pins on protein amino acid scale schematics represent experimentally determined cleavage sites and the heat maps illustrate the number of experimental cleavage sites per 50 AA window (darker red = more sites per window). c) receiver operating characteristic (ROC) curves evaluating experimentally detected MMP9 cleavage sites (by LC-MS/MS) from purified VTN and DCN proteins against PROSPER- and DeepCleave-predicted cleavage sites. Both algorithms struggled to accurately predict experimental sites in native proteins. d) MMP9-digested proteins detected by LC-MS/MS in a complex cell-derived matrix. A total of 60 proteins were detected from which 40 had at least one non-tryptic cleavage site (assumed to be due to MMP9). e) ROC curves evaluating experimentally detected MMP9 cleavage sites (by LC-MS/MS) from a complex cell derived matrix against PROSPER- and DeepCleave-predicted cleavage.

In a parallel set of experiments, we generated a complex ECM proteome by decellularizing a post-confluent culture of HDFs and digesting the deposited matrix with MMP9. In the baseline (non-MMP9 digested) matrix we identified over 60 proteins, many of which (COL6A1, COL6A2, COL6A3, EMILIN1, COL1A1, COL1A2) are found in human dermis (36,49). LC-MS/MS identified putative MMP9 cleavage sites in 40 out of 60 proteins including (COL6A1, EMILIN1, FBLN2, COL1A2) resulting in 390 putative MMP9 cleavage sites **(Fig.2.d)**. As with their performance against individual purified native proteins, both PROSPER and DeepCleave struggled to accurately predict cleavage sites in this a complex mixture of native ECM proteins (PROSPER AUC: 0.56; DeepCleave AUC: 0.6) **(Fig.2.e)**. This may be due to the experimental training data relevance: 1) different methods used in determining cleavage sites in our experiment vs MEROPS; 2) the quantity of the data available in such databases and/or; 3) the need for a more sophisticated algorithm. For example, all cleavage site identities were generated by these algorithms using MEROPS data which contains only 53 MMP9 human protein substrates corresponding to 301 cleavage sites derived from a mixture of experimental designs and tissues (27). These performance results highlight the need for better proteolysis models which utilise not only state-of-the-art machine learning approaches but also more expansive, up-to-date training datasets of native protein substrates, calibrated against experimental data.

### Protease cleavage site prediction performance can be improved using a deep bidirectional recurrent neural network architecture

Given the relatively poor performance of existing algorithms in predicting MMP9 cleavage in native ECM proteins we next aimed to develop and evaluate a new protease cleavage prediction algorithm. In order to better understand the complex proteolysis in a tissue such as ageing/inflamed skin, it is important that developed computational models are capable of accurately predicting cleavage sites for a wide range of proteases for which there is limited experimental evidence. Recurrent neural network (RNN) architectures, initially developed for natural language processing, are particularly suited to model genomic and proteomic sequences (59). This is particularly important for modelling proteolysis, as the AAs surrounding the cleavage sites play a significant role in determining the cleavage specificity (60). RNNs have recently achieved notable success in the field of proteomics, but have not yet been used to model proteolysis (35,61). Here we adapted the methodologies employed by both PROSPER and DeepCleave to develop a novel, deep bidirectional recurrent neural network (deep BRNN) based proteolysis prediction algorithm calibrated against inhouse experimental datasets. For data collection, the training, testing and validating datasets for serine proteases and MMPs were collected from the MEROPS database (27) and supplemented with additional MMP cleavage sites from a 2016 study (62) by Eckhard et al. Data were collected for two groups: MMPs (−1, −2, −3, −7, −8, −9, −12 and −13) and other proteases (Elastase 2, Granzyme B, Cathepsin -K, -G, -B, -D). These published datasets were also supplemented with the cleavage site information generated from our MMP9 LC/MS-MS experiments **(Fig.3.a)**.

**Figure 3.**
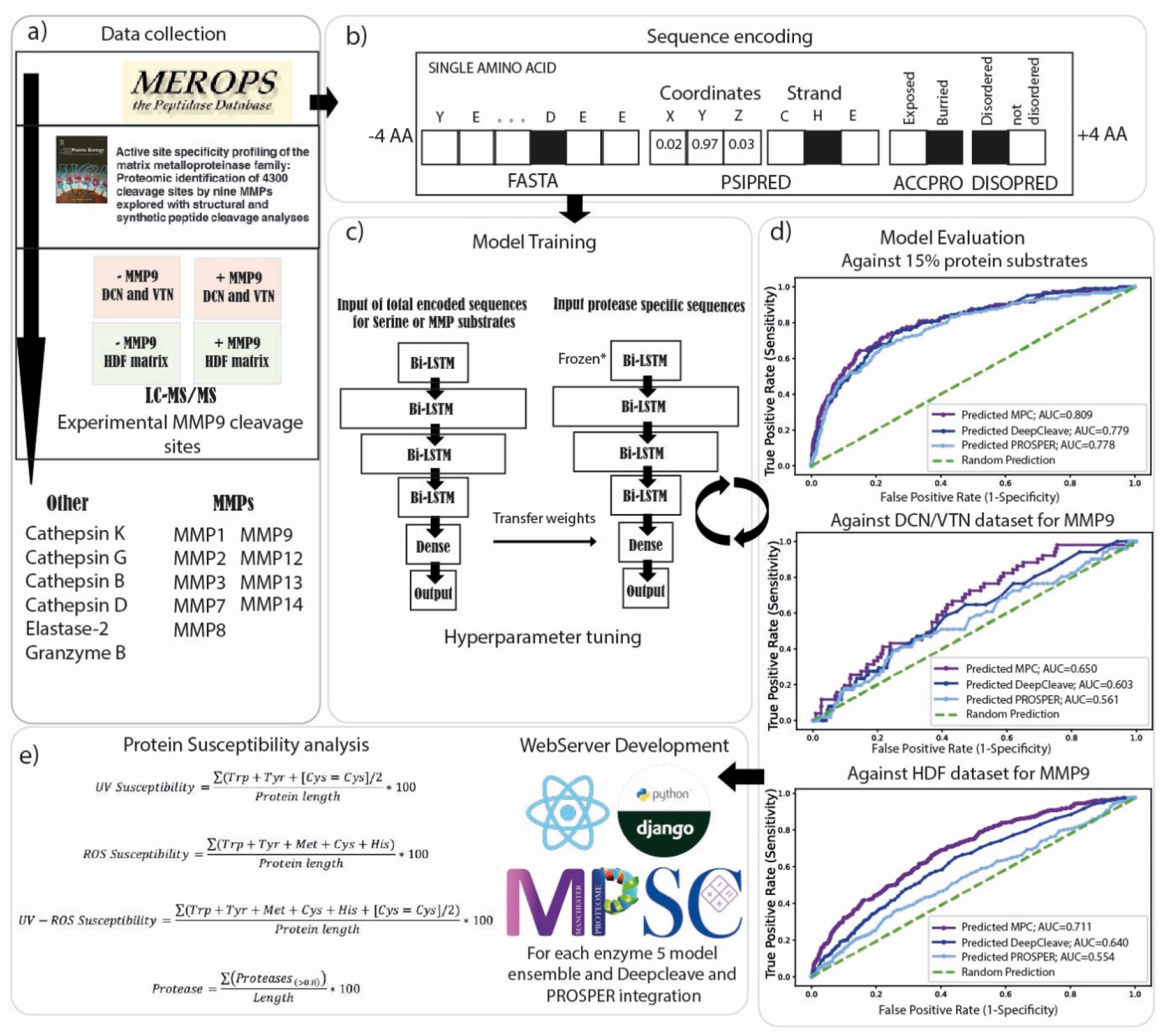
Manchester Protease Cleavage prediction model: architecture and testing. a) We have developed a novel protease prediction algorithm – termed Manchester Proteome Cleave (MPC) based on the state-of-the-art deep learning methodologies. The development of MPC involved five major steps: a) Data Collection: integration of data from publicly available protease cleavage databases (MEROPs), the Eckhard 2016 dataset and our own in-house experiments. b) Sequence encoding: MPC utilised primary AA sequence and structural features derived by in silico modelling with PSIPRED, DISOPRED and ACCPRO. One-hot encoding was used to represent the protein sequences in the format required for recurrent neural networks. c) Model training: the MPC model consisted of 4 bidirectional long-short term memory (Bi-LSTM) layers, one fully connected dense layer and an output layer. A general protease model was first pretrained and the resultant weights were transferred to train a protease-specific model. d) Model evaluation: Hyperparameters were tuned against 15% of excluded proteins and for MMP9 also against our experimental DCN/VTN and HDF cleavage site dataset utilising a grid search strategy. e) Webserver Development and Protein Susceptibility analysis: the best-performing architecture was used to train an additional four models which were integrated into our MPSC webtool. Hence MPSC used 5 model averaged ensembles to infer the final cleavage scores for each of the proteases. MPSC webtool is also capable in performing UV and ROS assessment of proteins. The MPC Webserver is publicly available on http://manchesterproteome.manchester.ac.uk/#/MPSC.

In addition to the primary AA sequences, secondary structure, disordered regions and solvent accessibility may also play significant roles in determining the probability of protease cleavage (30). We therefore used the methods previously employed by PROSPER (PSIPRED (63), DISOPRED2 (64) and ACCPRO 5.2 (65,66)) to predict these structural features for the whole human proteome prior to sequence encoding and model training. Each AA for every protein, in combination with these structural features, was encoded in a format suitable for RNNs. For sequence encoding, a sliding 8 AA window (4 AAs upstream and 4 AAs downstream of the predicted cleavage site) was utilised **(Fig.3.b)**. To build a cleavage site prediction model we needed to address two challenges: i) there is limited data available that reports MMP9 substrate specificity (e.g. MEROPs containing only 53 protein substrates corresponding to 301 cleavage sites), which can be used to develop deep learning models **(Supporting File 3)** and; ii) the large imbalance between cleavage sites vs non-cleavage (i.e. the number of non-cleavage sites vastly outweighs the number of cleavage sites) resulting in unbalanced datasets. The first challenge was addressed by: 1) complimenting the MEROPS cleavage data with data available in the Eckhard 2016 study (62) and; 2) using transfer learning approaches, where a general protease cleavage model was pretrained using all the available data (separately for MMPs and other proteases) and subsequently used as a starting point to train protease-specific models (using only the cleavage data for a specific protease) (67). The second issue was addressed by weighting the model prior to model training to “pay more attention” to the minority (cleavage sites) class. For model training, different architectures, depths and hyperparameters of deep learning networks were evaluated in terms of AUC, F1 and MCC scores against cleavage sites sourced from three test sets: i) protein substrate identities for each protease from the MEROPS and the Eckhard 2016 study which were excluded from the training datasets (consisting of 15% of the total number, randomly selected); ii) the DCN, VTN and; iii) HDF MMP9 cleavage datasets as previously described. The best performing architecture was composed of four bi-LSTM layers, one dense layer and one fully connected layer **(Fig.3.c)**. Crucially, the best performing setup, after adjusting various parameters, could more accurately predict cleavage sites in ECM proteins compared to PROSPER and DeepCleave for all test sets evaluated by AUC, F1 and MCC scores. The AUC scores for MMP9 for: i) the 15% protein test set were MPC: 0.809, DeepCleave: 0.779 and PROSPER: 0.778; ii) the DCN & VTN test set were MPC: 0.650, DeepCleave: 0.603 and PROSPER: 0.561 and; iii) the HDF data were MPC: 0.71, DeepCleave: 0.640 and PROSPER: 0.554 **(Fig.3.d)**, respectively. Similarly, MPC performed better for the 15% protein test set than PROSPER and DeepCleave for all other modelled proteases. For each of the proteases there was some agreement between MPC, DeepCleave and PROSPER predicted cleavage sites; however, each algorithm had also identified cleavage sites unique to the model **(Supporting File 3)**.

### The MPSC webtool can distinguish between protease susceptible and resistant proteins and reveal novel potential targets of proteolytic degradation distinct to UVR/ROS

Having developed both a model capable of predicting oxidative and/or UVR susceptibility and a new algorithm capable of predicting individual protease cleavage sites, we next aimed combine these into a webtool suited to predicting all these aspects of protein susceptibility to damage. In a similar approach to UV/ROS calculations, we calculated the number of protease cleavage sites per length of the protein which we correlated to protein susceptibility. To create a more encompassing statistical output with a better generalisation capability, the final output of MPSC consists of an ensemble model averaging five MPC model outputs **(Fig.3d)**. The Manchester Proteome Susceptibility calculator (MPSC) is available on http://manchesterproteome.manchester.ac.uk/#/MPSC (or http://130.88.96.141/#/MPSC) where users can perform their own analyses of relative protein susceptibility to UVR/ROS and selected proteases.

The HDF-derived matrix experiment not only provided information on MMP9 substrates but also revealed 20 proteins which did not appear to have LC-MS/MS-detectable MMP9 cleavage sites. These 20 proteins included COL3A1, COL12A1, LTBP2, VCAN and others **(Fig.2.d)**. We have used the identities of these cleaved and non-cleaved proteins to determine whether predicted MMP9 protease susceptibilities differed statistically between these two groups. The MPSC score for MMP9 was set to 0.8 which corresponds to highly confident cleavage sites. Using this approach, proteins with no experimentally detected MMP9 cleavage in the HDF-derived proteome also had a significantly lower predicted MMP9 susceptibility than the proteins that had at least one MMP9 cleavage site (*p* = 0.005, student’s *t* test). This analysis suggests that novel substrates for proteolytic degradation may be identified using *in silico* proteolytic modelling.

We next applied MPSC webtool calculations to every ECM and extracellular protein in the Manchester Skin Proteome (36) in order to predict their susceptibility to UVR/ROS and to UV-upregulated skin proteases (interstitial collagenase (MMP1), 92kDa gelatinase (MMP9), stromelysin-1 (MMP3), Cathepsin K, and Granzyme B) (55,68).

Using the MPSC webtool, we initially analysed skin ECM proteins for the predicted protease and UVR/ROS susceptibilities using MPSC-MPC models. This analysis suggests that tropoelastin (the elastin precursor) will be particularly susceptible to MMP-mediated cleavage which agrees well with experimental observations (69). In addition, multiple proteins involved in elastic-fibre formation and organisation (FBLN1, FBLN2, FBLN5 EFEMP1, EFEMP2, MFAP4 and FBN2) were also predicted to be susceptible to not only MMP proteolysis but also to UVR/ROS. These predictions suggest that both protease mediated and UVR/oxidative damage may play a role in the deterioration of the elastic fibre architecture which is characteristic of photodamage in human skin (70). We have previously shown an interplay between these mechanisms whereby UVR exposure enhances the degradation of fibrillin microfibrils (52).

In addition to elastic fibres, collagens also undergo profound remodelling in photo-exposed skin. Protease-mediated activity has long been suspected as a driver of age-related degradation of skin collagens (44,45) but given the intermittent and/or chronic low-level action of these mechanisms, the causative link has yet to be established. Whilst the protease susceptibility of these major collagens has been confirmed experimentally (71), our computational techniques also suggest that many skin collagens (i.e. II, III, IV, V, VI, XII, and XV) will be degraded by proteases (MMP −1, −3, −9, Granzyme-B, and Cathepsin K) but not by UVR/ROS. For the ubiquitous microfibrillar collagen VI assemblies, protein abundance and architecture is resistant to photodamage (16). The *in vivo* resistance of collagen VI to photodamage (16) suggests that this ECM assembly is resistant to multiple degradative agents. However, the alpha 5 chain of COL6 is degraded by Cathepsin K (72,73) and we have shown subtle UV-induced changes in the structure-associated features by mass spectrometry (15). We therefore conjecture that differential mechanisms may drive the degradation of elastic fibre components and skin collagens, necessitating different preventative and/or therapeutic approaches.

In addition to predicting the relative susceptibility of major structural ECM assemblies, our analysis also suggests that protease-mediated proteolysis of basement membrane proteins including laminins (LAMB-1, −2, −3, −4, LAMA-3, −4) and nidogen (NID1) may be a contributing factor to the remodelling of the dermal-epidermal junction in photoexposed skin (74). Other proteins predicted to be highly susceptible to MMP degradation include several proteoglycans (OGE, ASPN, KERA, and DPT), retinoic acid receptor responder protein 2 (RARRES2), matrix Gla protein (MGP), procollagen C proteinase enhancer (PCOLCE), TGFβ1, apolipoprotein E (APOE) and Von Willebrand Factor (VWA5B1). Some proteins appear to be susceptible to only a single enzyme, such as collagen triple helix repeat-containing protein 1 (CTHRC1: Granzyme B) and adiponectin receptor protein 1 (ADPOQ: cathepsin K) **(Fig.4.b-f)**.

**Figure 4.**
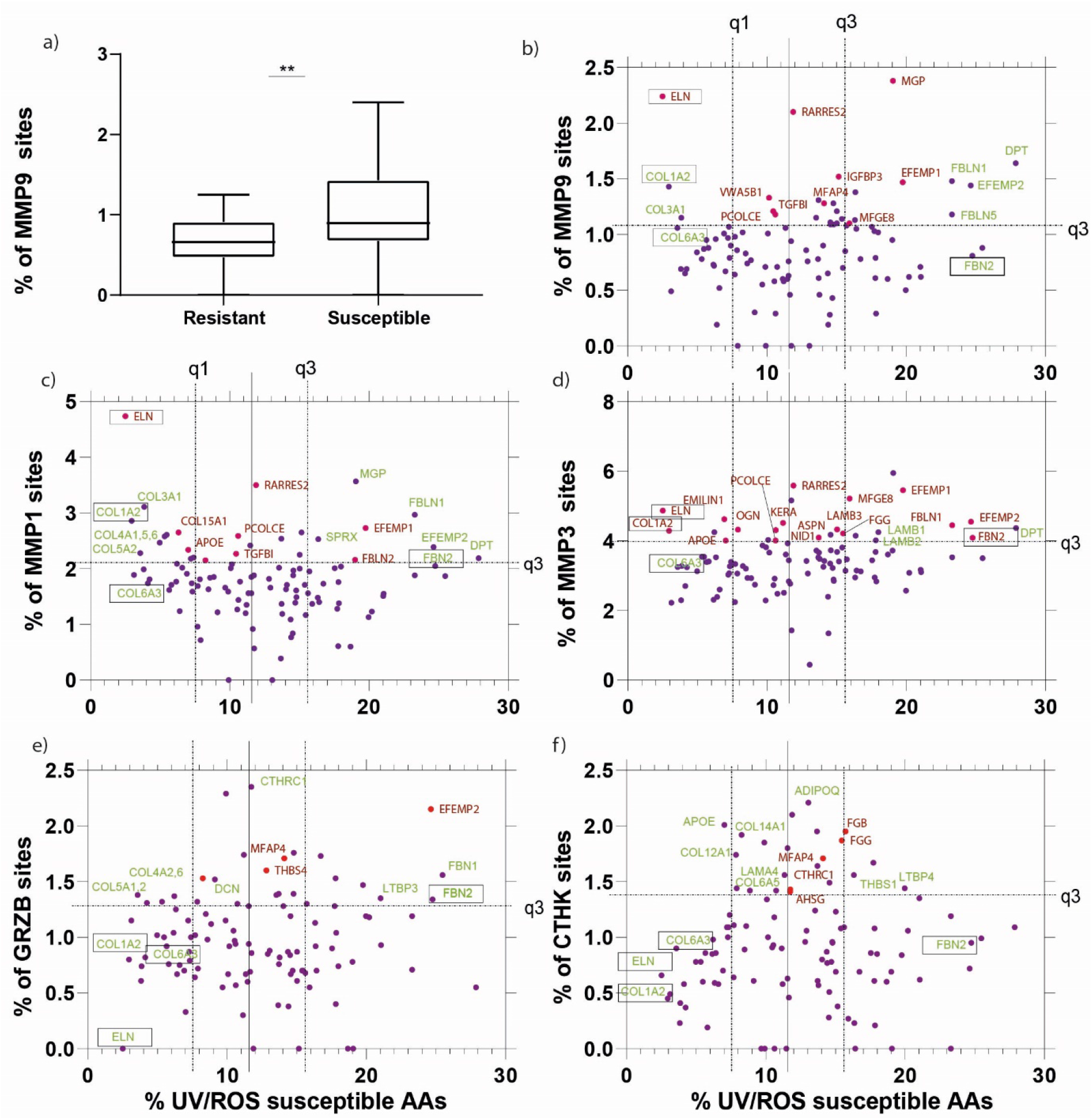
UVR/ROS vs protease susceptibilities of MMP-1, 3 and 7, Granzyme B and Cathepsin K of skin proteome-derived ECM components. a) Using the protein identities in the MMP9 degradation dataset, which had no MMP9 cleavage sites or had at least one cleavage site, allowed the evaluation of MPC model’s performance to predict novel substrates. Proteins that had no experimental cleavage sites yielded significantly lower predicted MMP9 susceptibility scores. Having defined that susceptibilities could be analysed using the MPC model, we used these models and analysed the susceptibilities for MMP– 1 (c), 3 (d), and 9 (b), Granzyme B (e), and Cathepsin K (f). Proteins which were predicted as highly susceptible by PROSPER or DeepCleave only are labelled in red text. Proteins that are predicted as highly susceptible by MPC only are labelled in green text. Throughout, we have highlighted (boxed text) important skin proteins COL1A2, FBN2, ELN and COL6A3 which merit discussion.

In addition to structural ECM components, photoageing may also affect non-structural extracellular proteins such as growth factors, antioxidants and enzymes which play key roles in the maintenance and regeneration of normal healthy skin. Our analysis suggested that MMPs, Cathepsin-K and Granzyme-B are capable of digesting (or self-digesting) each other in addition to other catalytic proteins and inhibitors; such as Cathepsins themselves (A, C, K, H, B, Z, V), Kallikreins (KLK5, KLK6, KLK7, KLK13), MMPs (MMP10, MMP11, MMP19, CPA3), PLAU, ADAMTS17, CFI, and protease inhibitors TIMP1 and 2, SPINK6, CST6, PI3, SERPINF1, SERPINA1, 3, 12, SERPINB5. It has been shown previously that proteases exhibit complex synergetic relationship, cleaving one another to amplify protease cascades such as those mediated by the so-called “cysteine switch”. (75). This consistent prediction of such catalytic proteins being protease susceptible markers and the extensive literature of protease activation mechanisms strengthens our confidence in the predictions of the MPC. Defensin family members (DEFA1B, DEFB1, DEFB4B) - which play antimicrobial roles in skin - are predicted by the MSPC to be highly susceptible to both UVR/ROS and proteases, thereby suggesting their role not only as antimicrobial peptides but also potential sacrificial sunscreen-like proteins and antioxidants (46,76). Other examples of proteins predicted to be susceptible to both proteolysis and UV/ROS include IGF1, TGFα, INHBA, NTF3, VEGFC, HGFAC, TSHB, and SLURP1. Markers predicted to be susceptible to proteolysis but resistant to UVR/ROS include Galectins (LGALS1, LGALS7B, LGALSL) and Annexins-1, 3, 5 (ANXA1, ANXA3, ANXA5) potentially indicating their need to be resistant to environmental factors **(Fig.5.a-e)**.

**Figure 5.**
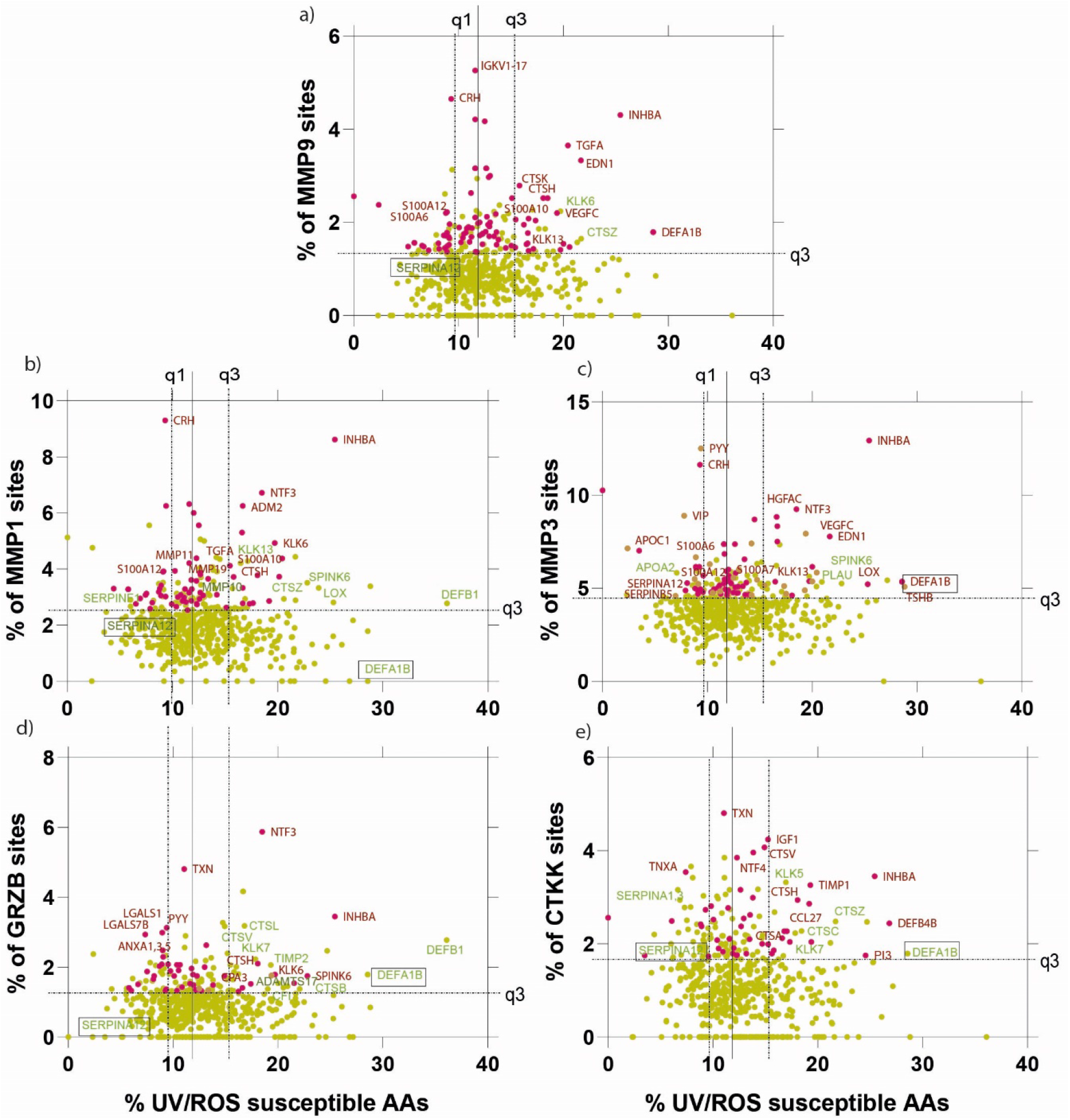
UVR/ROS vs protease susceptibilities of MMP-1, 3 and 7, Granzyme B and Cathepsin K of skin proteome-derived extracellular components. Similarly to the ECM components we have also investigated proteins in the extracellular space important in maintaining the ECM. We analysed extracellular components for MMP-1 (b), 3 (c), 9 (a), Granzyme B (d) and Cathepsin K (e). Proteins which were predicted as highly susceptible by PROSPER or DeepCleave only are labelled in red text. Proteins that are predicted as highly susceptible by MPC only are labelled in green text. Throughout, we have highlighted (boxed text) important skin proteins which merit discussion.

## Discussion

Degradomics is an ever-expanding field with the potential to impact translational research by deepening our understanding of tissue regeneration processes as well as contributing to drug discovery efforts (9,77). In this work we show that analysis of AA sequence and extracted structural features, in combination with state-of-the-art deep BRNN, is capable of predicting proteolytic cleavage sites with a better degree of accuracy.

Furthermore, we can use these predictions and sequence chromophore counts to identify not only known protein targets of photodamage but also potential novel protein targets and the relative susceptibilities within key protein families. With this work, we have stratified proteins within the entire human skin proteome to reveal their predicted susceptibilities to UV, ROS and proteolysis providing significant novel insights in skin research. Particularly, we have highlighted the fact that fibrillar collagens are predicted to be preferentially degraded by proteases alone whereas elastic fibres are likely to be susceptible to UVR/ROS and proteases in combination. We also demonstrate that proteins involved in catalytic processes have a high percentage of predicted confident cleavage sites per protein length. Moreover, we have made this analysis applicable to any AA sequence of interest. Beyond the field of dermatology, the concordance between our computational analyses and previously reported observations of both *in vitro* and *in vivo* protein degradation suggests that this approach has the potential to identify novel protein biomarkers for tissues subjected to inflammation or ageing-related disease.

Whilst the MPC model performed better than existing models, its success in predicting experimental cleavage sites in native proteins still requires further improvement. Furthermore, while we have attempted initial experimental MMP9 cleavage site detection in a complex HDF proteome using LC-MS/MS approaches, it is clear that many cleavage sites may have been missed, particularly for low abundance proteins, where the peptide-coverage of the protein is also low; therefore, experimental approaches and sample preparation methods require even further improvement. The difficulty in predicting cleavage sites in native proteins may be partly due to the heterogeneous nature of the data currently available in public databases such as MEROPS upon which the algorithms were built. For example, elastin degradation by proteases is very well-defined, but well-known MMP substrates (such as FBN1) are not been necessarily reflected in the available databases (27,69,78). Also data referenced in these databases is drawn from multiple experimental methods applied to disparate biological samples including peptide libraries, extracted cartilage proteins, *post mortem* brain tissue, and many others which may not necessarily translate to a different *in vitro* systems (27). Critically, we have shown using an LC-MS/MS peptide location fingerprinting approach that ECM proteins can exhibit tissue-dependent structures (79). Furthermore, higher order structures play a crucial roles in determining proteolysis; however, currently the accurate prediction of protein secondary and tertiary features remains challenging (80). In this context, improved algorithms trained on expanded experimental datasets with additional informative features would benefit proteolytic cleavage site prediction. Future development of these techniques will depend on the availability of training data encompassing key target proteomes and proteases and improved prediction by taking into consideration the impact of local protein structures on the protease-specific cleavage outcomes.

Overall, MPSC webtool builds upon the success of the Prosper and DeepCleave algorithms which have contributed significantly to the degradomics fields and can further assist in novel biomarker discovery, reveal the primary AA sequence susceptibilities to UV/ROS and proteases and even assist in a novel bioactive matrikine discovery (81).

## Materials and Methods

### Dataset collection

FASTA sequences and disulphide bonds for each protein in the Manchester Skin Proteome (MSP) were retrieved from the Uniprot database in Jan 2020 (82). For MPSC development, experimentally validated protein substrate annotations were collected from Eckhard 2016 and MEROPS (v12.2) in Jan 2020 (27,62). Only human protein cleavage sites were retrieved. The entire MSP (36) and proteins present in the Eckhard 2016 were digested using locally installed PROSPER and full proteome digestion by DeepCleave was provided by our Monash collaborators.

### Deep RNN protease model

#### Evaluation Metrics

As the dataset is highly imbalanced containing thousands of non-cleavage sites vs only hundreds of cleavage sites, and thus in order to evaluate the performance of PROSPER, DeepCleave and MPC, we used F1 score representing the harmonic mean of precision and recall; the Matthews’ Correlation Coefficient (MCC), Precision, Recall and Area Under the Curve (AUC) scores. Performance differences of classifiers are visualised with receiver operating characteristic (ROC) curves (sensitivity - [true positive rate] against the 1-specificity [false positive rate] and evaluated with area under the curve (AUC) score. MCC and F1 score are defined as follows:

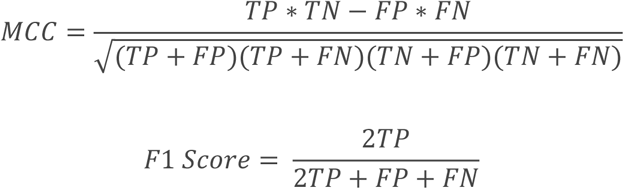

where *TP* represents the number of true positives, *TN* represents the number of true negatives, *FN* represents the number of false negatives, and *FP* represents the number of false positives, respectively.

#### Feature extraction

PROSPER revealed that not only the AA sequence context surrounding cleavage sites, but also protein secondary structure, disordered regions and solvent accessibility play important roles in proteolytic cleavage site prediction (30). The whole human proteome was annotated with PSIPRED (63), DISOPRED2 (64) and ACCPRO 5.2 (65,66) to retrieve these different types of sequence-derived structural features.

#### One-Hot encoding

Each AA was encoded using the one-hot encoding, resulting in a 20-dimenisonal vector where each dimension represents one of the 20 common AAs. To gain the insights from the surrounding AAs a sliding window of −4 and +4 AAs additional to the cleavage site was used for each sequence. At the N- and C-termini where there are no following AAs, each position of 20-dimensional vector contained only zeros. Furthermore, we complemented each of these vectors with three-dimensional coordinates retrieved from SCRATCH (x, y, z), one-hot encoded representation of whether amino acid was part of coil, strand or helix, two-dimensional one-hot vector encoding whether it was exposed or buried, as well as two-dimensional one-hot vector of whether AA was disordered or not.

#### Train, test and validate data split

In contrast with previously published models, which split the data in train, test and validate datasets on the cleavage site basis regardless of the protein source, we have ensured that the testing, training and validating data all come from independent proteins. We used TensorFlow random seed to ensure the consistency of evaluations and ensured that the same protein identities were selected each time when evaluating the model performance; 70% of the proteins were used for the training, 15% for testing and 15% for validating.

#### Architecture of the Deep RNN

We used the Python 3.8 TensorFlow Keras package to implement our MPC model. There are many more non-cleavage sites than the cleavage sites resulting in a very imbalanced dataset. To overcome this, we assigned larger weights to cleavage sites than non-cleavage sites, enforcing the classifier to “pay more attention” to the underrepresented class. We also utilised transfer learning to overcome the issue with limited amounts of data available for certain proteases (67). We ensured that the general model and protease-specific model contained the same identities for testing, training, and validating. MPC protease models consists of four bidirectional LSTM layers, fully connected Dense layer and an output layer. We set epochs to a very large number (>10 000) and monitored the early stopping by measuring the maximum F1 score with patience of 20 epochs. A generic model utilised all the MMPs or other proteases. We then used transfer learning to convert the generic model into protease-specific model by loading the initial weights, freezing the initial layer and continuing the training process with the data that only had protease specific entries. Our experimental DCN/VTN and HDF MMP9 datasets were used to tune the hyperparameters (the number of AAs surrounding cleavage site, learning rate, batch size etc.) to achieve the optimal performance in the skin-relevant model system. Subsequently, based on the optimised hyperparameters we trained 5 models for each of the proteases and took the average output score as the final prediction.

### UVR/ROS and protease MPSC susceptibility model calculations

Previous analysis has revealed clear agreement between experimental UV and ROS susceptibility and chromophore content of the FASTA sequence. To implement the UV, ROS and combined UVR/ROS model, simple mathematical equations were derived which quantify the number of UV and ROS chromophore AAs per length of the sequence. For proteolysis susceptibility calculations highly confident (score > 0.8), MPC cleavage sites were selected and the number of cleavage sites per protein length calculated. The calculations employed in MPSC are:

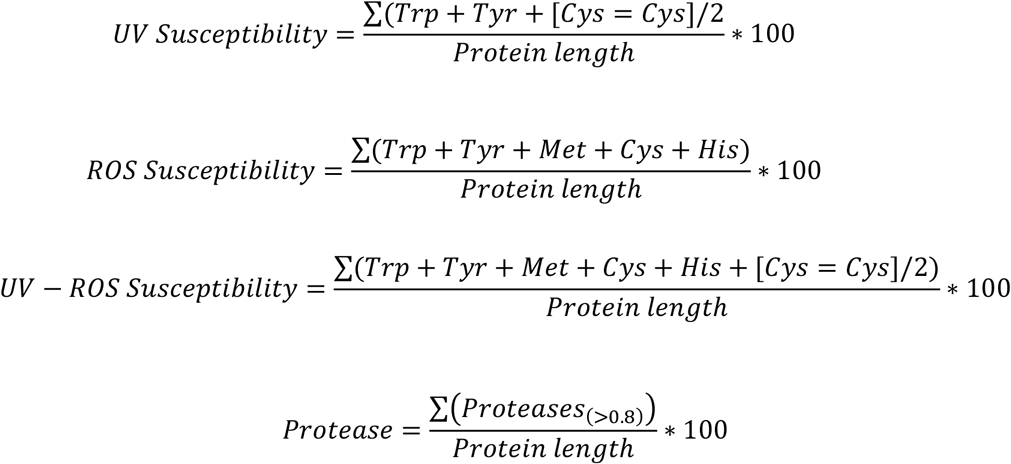

where Trp represents tryptophan, Tyr represents tyrosine, Met represents methionine, Cys represents cysteine, His represents histidine and Cys=Cys represents disulphide bound cystine.

### LC-MS/MS methods for generating experimental testing dataset

#### Cell culture

Human dermal fibroblasts (HDFs) were cultured from a scalp biopsy, obtained from a hair transplant procedure. Ethical approval was obtained from the National Research Ethics Committee (ref 19/NW/0082) and the tissue donor provided written, informed consent prior to the procedure. Skin biopsies were cultured dermis-side down in Dulbecco’s modified Eagle’s medium (DMEM) (Thermo Fisher) containing 20% (v/v) foetal bovine serum (FBS), GlutaMAX (2 mM), penicillin (100 U/mL), streptomycin (0.1 mg/mL) and amphotericin-B (2.5 μg/mL), until HDFs were observed migrating out of the explant. Culture medium was changed every 2-3 days until cells reached 80% confluence (approx. 3 weeks), at which point explanted tissue was aseptically removed from the culture and HDFs were sub-cultured using trypsin (0.05% w/v) - ethylenediaminetetraacetic acid (EDTA; 0.02% w/v). Subsequent culture of HDFs used supplemented DMEM containing 10% (v/v) FBS. For extracellular matrix (ECM) deposition, HDFs at passage 4 were seeded at 1.2X10^5^ cells/mL, in 0.5 mL medium containing 50 μg/mL ascorbic acid (final concentration), in a 24-well plate, and cultured until confluence (approx. 2 days). At this point, 0.25 mL medium was removed and replaced with fresh medium. Medium was replaced every 2-3 days.

#### *HDF-deposited ECM* in vitro

The procedure for obtaining HDF-deposited ECM was carried-out as previously described (83). Briefly, medium was removed from cultures on day 9 post-confluency and the cell layer washed twice with phosphate buffered saline (PBS). To remove cells, 0.5 mL of extraction buffer (20 mM ammonium hydroxide solution, 0.5% v/v triton-X 100, in PBS without Ca^2+^/Mg^2+^) was added to HDFs for 5 min and cell depletion was determined by repeatedly checking cultures under a light microscope. Extraction buffer was diluted by addition of PBS and HDF-depleted ECM was incubated at 4°C overnight. Following removal of diluted extraction buffer, the remaining cell debris was removed by three successive washes in PBS. The presence of deposited, HDF-depleted ECM was determined by light microscopy.

#### *MMP9 degradation of ECM and DCN/VTN* in vitro

Human recombinant MMP9 (hrMMP9; R&D Systems) was activated by adding 1 mM (final concentration) ρ-aminophenylmercuric acetate (APMA) and incubating for 24h at 37°C. HrMMP9 activity was confirmed by measuring cleavage of the fluorogenic peptide, Mca-PLGL-Dpa-AR-NH2 (R&D Systems), according to manufacturer’s instructions. HDF-depleted ECM was incubated with 10 μg/mL of active hrMMP9 whereas DCN (ab167743, abcam) and VTN (ab94369, abcam) was incubated at a 1:2 protein to active hrMMP9 ratio, diluted in MMP buffer (50 mM Tris, 150 mM NaCl, 10 mM CaCl2 and 0.05% (w/v) Brij-35, pH 7.5) at 37°C with agitation, for 24h. For HDF ECM assay buffer was removed and dialysed for 4 h against ddH2O (Thermo Fisher) and samples freeze-dried overnight. DCN and VTN samples were ran on 4-12% NuPAGE Novex Bis-Tris Gels (1 μg of protein per well) using standard SDS PAGE procedures. Non-digested control bands and digested bands were excised using a scalpel blade. ECM or DCN and VTN incubated with MMP buffer alone served as protease negative controls respectively.

#### HDF laid down-matrix sample preparation

Each sample’s aliquot was added to SMART Digest™ trypsin (Thermo Fisher) beads and shaken at 1400 rpm overnight at 37°C. The next day, reduction and alkylation was performed. Next, SMART Digest Beads were removed using TELOS MicroPlate™ filter tips (Kinesis; Cheshire, UK) into 1.5 ml Eppendorf^®^ LoBind tubes (Eppendorf; Stevenage, UK) to minimise electrostatic loss of peptides and acidified with 5 μl of 10 % formic acid (FA). Biphasic extraction was performed to partition organic molecules and surfactants. To do this, 200 μl of ethyl acetate (EA; Sigma) was added to each sample in Eppendorf^®^ LoBind tubes and vortexed for 1 min. Samples were then centrifuged to ensure phases were separated and the upper layer removed using gel-loading pipette tips. These steps were repeated twice with a further 200 μl of EA and then the resultant aqueous bottom phase was vacuum dried for 2 hours in room temperature. Once dry, 200 μl of injection solution (5% (v/v) acetonitrile, 0.1% formic acid) was added to the tube and peptide desalting performed. For all desalting of peptide preparations, OLIGO™ R3 Reversed Phase Resin (Thermo Fisher) beads were used. Beads were wetted to be activated using 50 μl OLIGO™ R3 beads plus 50 μl of wet solution (50% acetonitrile in ultrapure water; 1:1). Twenty (20) μl of this mix were placed in detached TELOS filter tips and liquid was pipetted out using a standard p1000 Gilson pipette (Gilson; Middleton, Wisconsin, USA) and process repeated twice more. Twice beads were washed by adding 50 μl of wash solution (0.1 % FA in ultrapure water) and liquid pipetted out. Once the beads were activated 100 μl of sample was gently added to TELOS filter tips containing activated OLIGO R3 beads, allowing the peptides from the sample to stick to the beads. Liquid is pipetted out and the ‘flow through’ retained. The process was repeated with the remaining sample. Twice beads with bound peptides were gently re-suspended in 50μl of wash solution and the flow through discarded. Twice, beads were gently suspended in 50 μl of elute solution (50 % (v/v) acetonitrile, 0.1% formic acid) to release the peptides from the beads and liquid was pipetted out and retained. The collected eluted sample contained the peptides for analysis. Eluted samples were transferred to mass spectrometry vials and vacuum dried using Speed Vac (Heto-Holten; Frederiksborg, Denmark), ensuring peptide stability until mass spectrometer was available.

#### DCN and VTN gel sample preparation

Excised bands were first placed into a perforated well plate, and then shrunk by washing in acetonitrile for five minutes. To remove the acetonitrile from the samples these were centrifuged for one minute at 1500 rpm. Sample gel pieces were then dried in a vacuum centrifuge for 15 minutes. Next, samples were covered completely by 10 mM dithiothreitol (DTT) in 25 mM ammonium bicarbonate and incubated at 56°C for one hour to reduce the proteins. Samples were cooled to room temperature, then centrifuged to remove the DTT. Iodoacetamide (55 mM) in 25 mM ammonium bicarbonate was added, and the samples were incubated for 45 minutes in the dark at room temperature. The iodoacetamide solution was removed by centrifugation, and the samples were washed with 25 mM ammonium bicarbonate for 10 minutes. Samples were then washed with acetonitrile, followed by another wash with ammonium bicarbonate, then a final wash with acetonitrile, centrifuging in between steps to remove the previous wash. The samples were then centrifuged again, then dried out in a vacuum centrifuge. Next, 5 μl of a 12.5 ng/μl solution of trypsin along with 45 μl of a 25 mM ammonium bicarbonate solution was added to the dried samples. Samples were kept at 4°C for 45 minutes for samples to absorb the buffer. After this time, samples were covered completely in 25 mM ammonium bicarbonate. Samples were incubated at 37°C overnight. The following day, peptides were extracted by incubating with 20 mM ammonium carbonate for 20 minutes, then centrifuging into a fresh storage plate. Peptides were extracted a further two times (20 minutes each) using 5% formic acid in 50% acetonitrile, using the same storage plate to pool all the supernatant into the same tube per sample. samples were transferred to mass spectrometry vials and vacuum dried using Speed Vac (Heto-Holten; Frederiksborg, Denmark).

#### Peptide preparation for mass spectrometry

For LC-MS\MS analysis, 20 μl of Injection Solution (5% acetonitrile (ACN) + 0.1% formic acid (FA) in ultrapure water) was added to each dried peptide samples. Peptide concentration in solution was measured using Direct Detect Infrared Spectrometer (Millipore). According to the concentration detected for each sample, they were diluted to have 12 μl of solution with a concentration of 800 ng/μl, and samples were submitted to the Biological Mass Spectrometry Core Facility (University of Manchester).

#### Liquid chromatography tandem mass spectrometry

Mass spectrometry was performed according to the Facility’s protocols (84,85). Digested samples were analysed by LC-MS/MS using an UltiMate^®^ 3000 Rapid Separation LC (RSLC, Dionex Corporation, Sunnyvale, CA) coupled to a Q Exactive HF (Thermo Fisher Scientific, Waltham, MA) mass spectrometer. Peptide mixtures were separated using a multistep gradient from 95% A (0.1% FA in water) and 5% B (0.1% FA in acetonitrile) to 7% B at 1 min, 18% B at 58 min, 27% B in 72 minutes and 60% B at 74 minutes at 300 nl min-1, using a 75 mm x 250 μm i.d. 1.7 μM CSH C18, analytical column (Waters). Peptides were selected for fragmentation automatically by data dependent analysis.

#### Webserver development and data visualisation

MPSC webtool was developed using the ReactJS framework for the front end and Python Django as the backend for the API development. Data powering the MPSC webtool was stored on the Apache server on the mySQL database. Data for this publication were visualised using Graphpad Prism 8, Adobe Illustrator and python matplotlib libraries.

## Author Summary

MO has developed all the models, performed bioinformatic analysis, designed webtool and backend databases and algorithms, analysed all the data and wrote the manuscript. MO conceptualised the application of MPSC. AE and MO together performed the native protein experiments. AE reviewed the literature of the existing protein degradation. MJS, SAH and AE contributed to the conception and design of the study, the interpretation of results. MJS and AE contributed to the preparation of figures and to writing. JS, JR and FL helped to initialize the project and provided advice and expertise in field.

CP and CMG designed and performed MMP9 HDF experiments. CEMG and REBW and contributed to the interpretation of results. All authors contributed to reviewing and editing of the paper.

## Acknowledgements

We would like to acknowledge the help by the Biological Mass Spectrometry Core Facility and research IT HPC cloud computing facilities in University of Manchester.

