## Supporting File 2 for "Predicting and validating protein degradation in proteomes using deep learning"

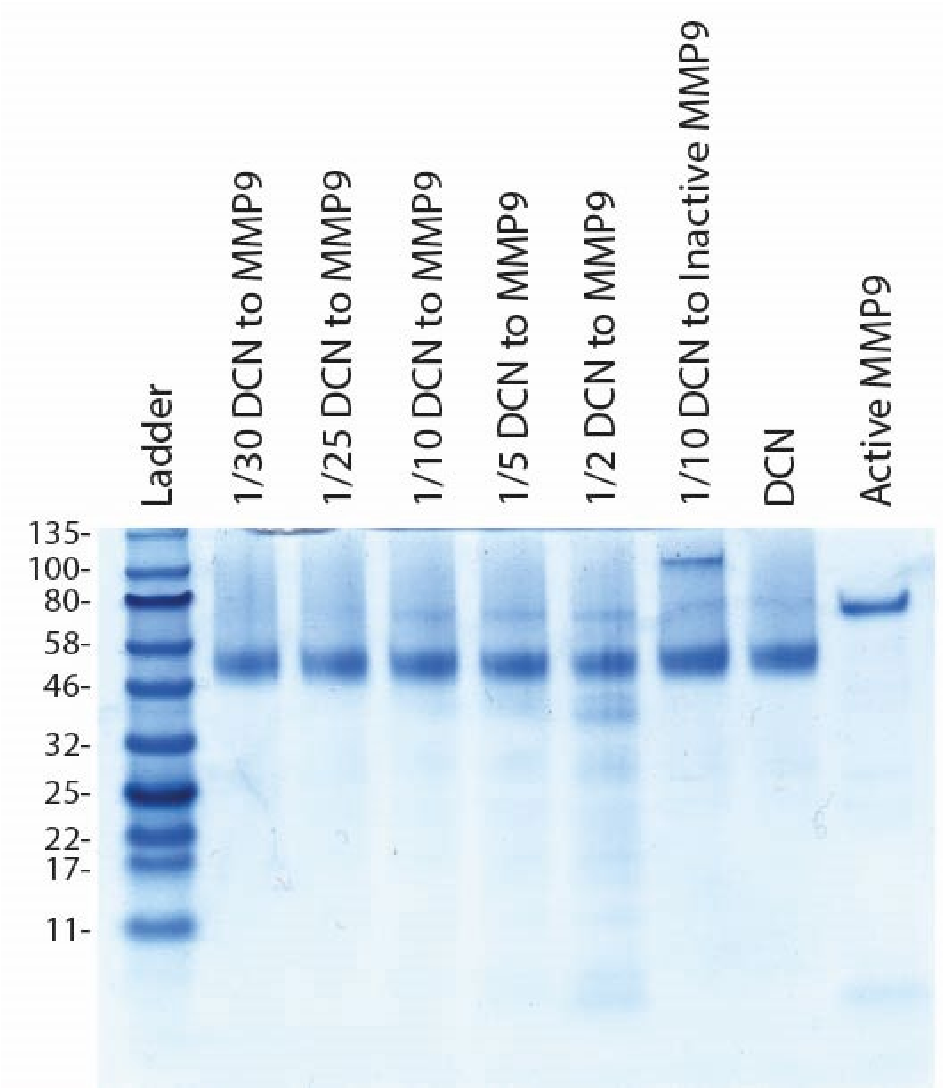


**Supporting Figure 1: Optimisation of protein to enzyme ratio.** MMP9 degraded DCN in a dose-dependent manor. At the highest DCN to MMP9 (1:2) multiple degradation products were readily detectable by SDS-PAGE. This relatively high enzyme concentration was used for subsequent experiments.

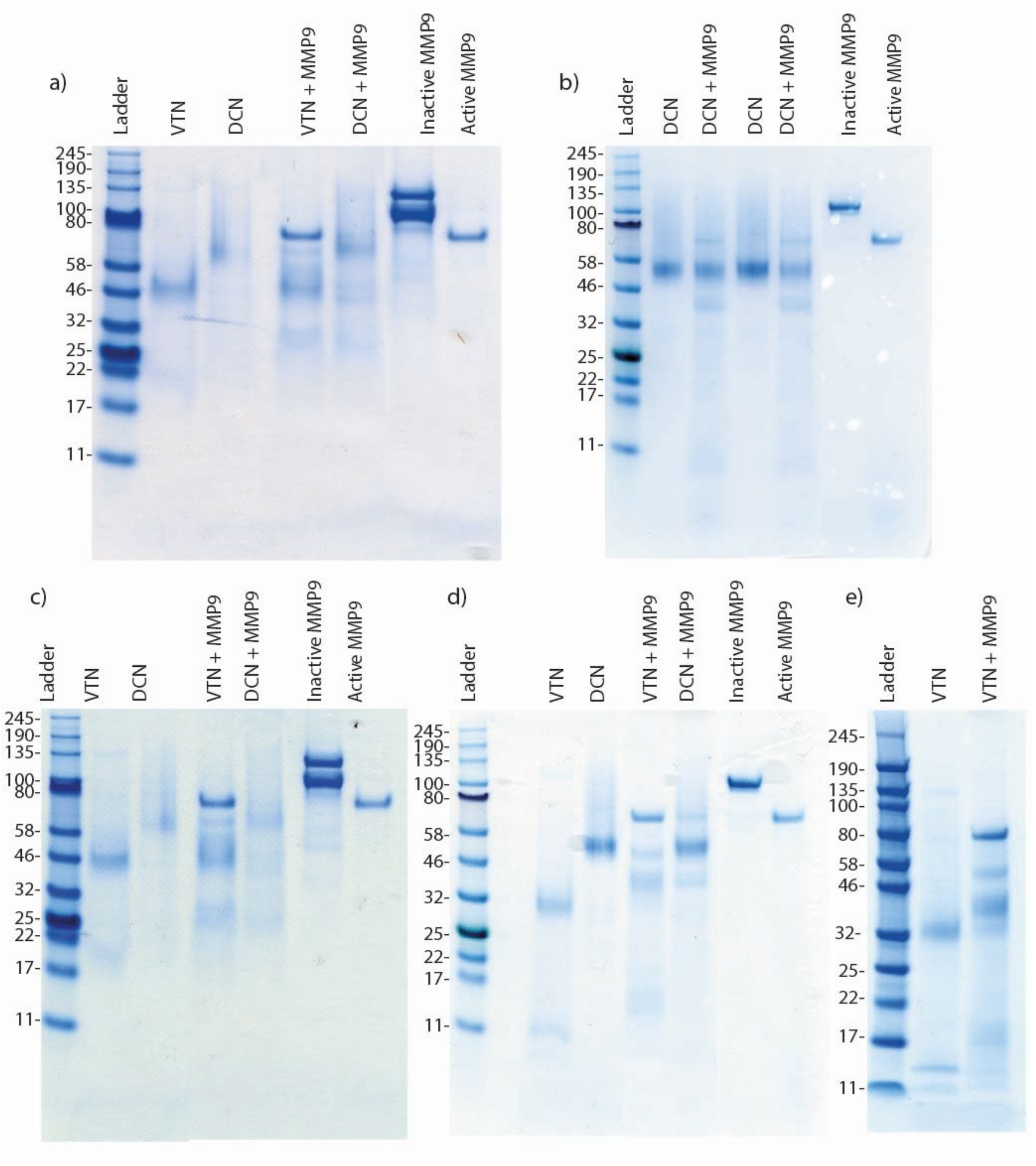


**Supporting Figure 2. Protein degradation experiments were performed in at least quadruplicate.** To confirm both the VTN and DCN degradation by MMP9 we ensured that both VTN and DCN experiments are performed at least 4 times. Five gels are showed here (a, b, c, d, e) which confirm the experimental consistency.
