## Supporting File 4 for "Predicting and validating protein degradation in proteomes using deep learning"

### MMP1 model performance against 15% test data

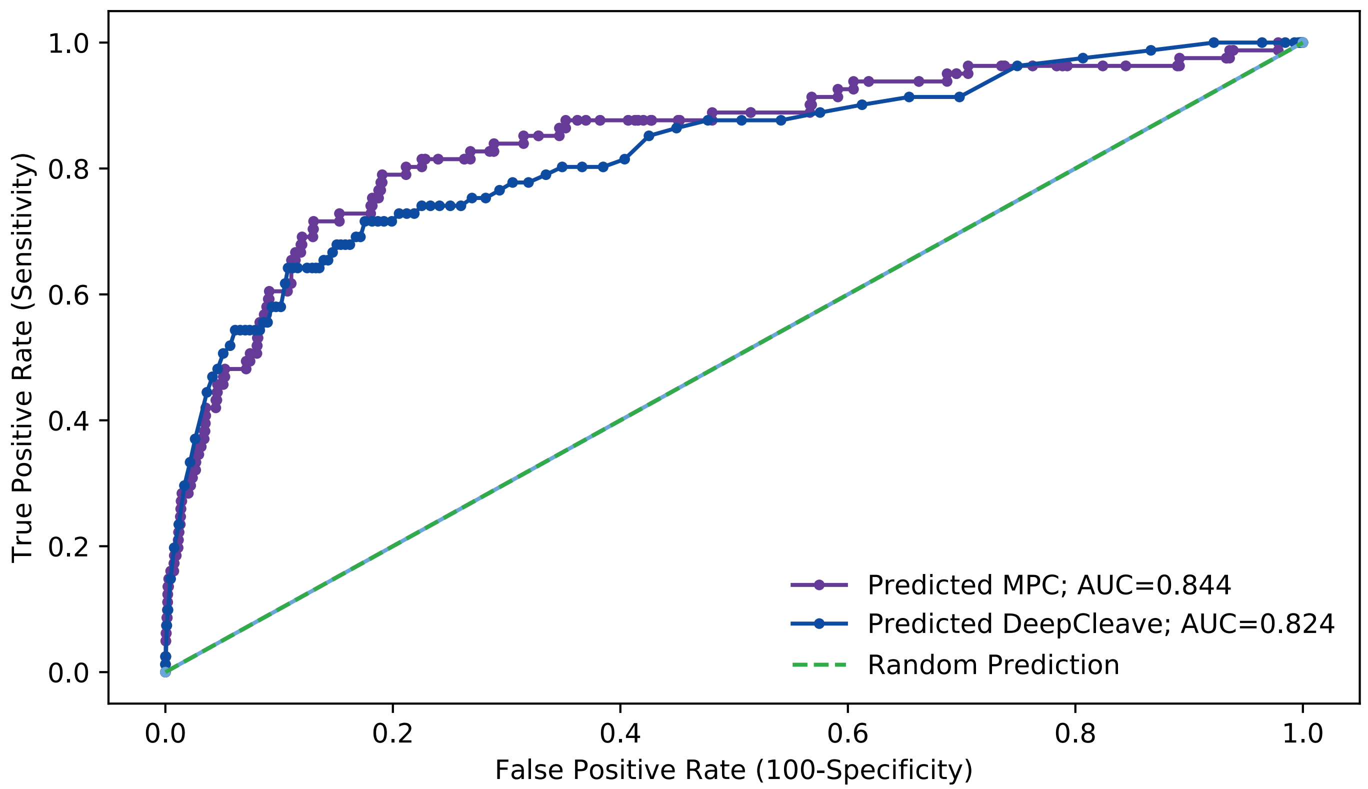


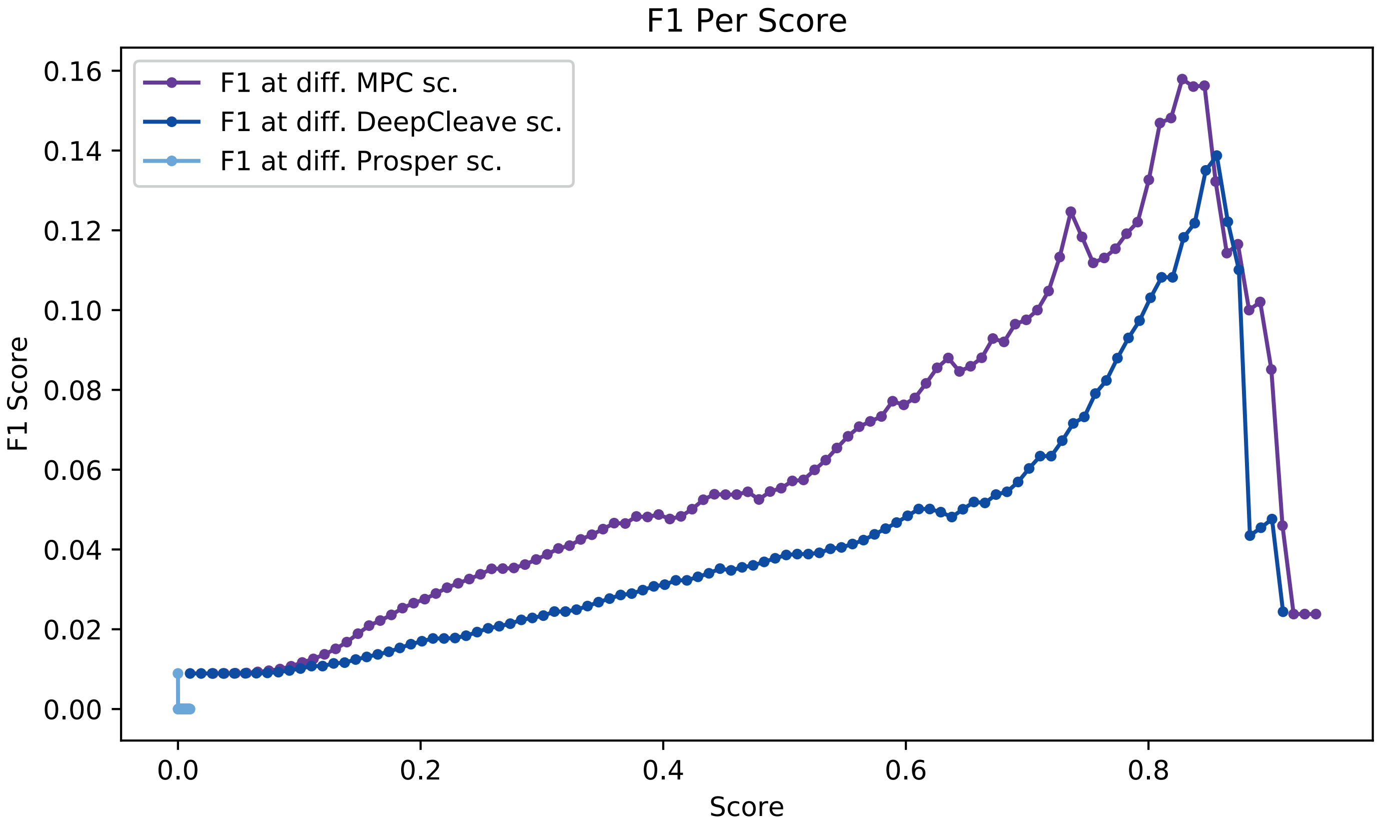


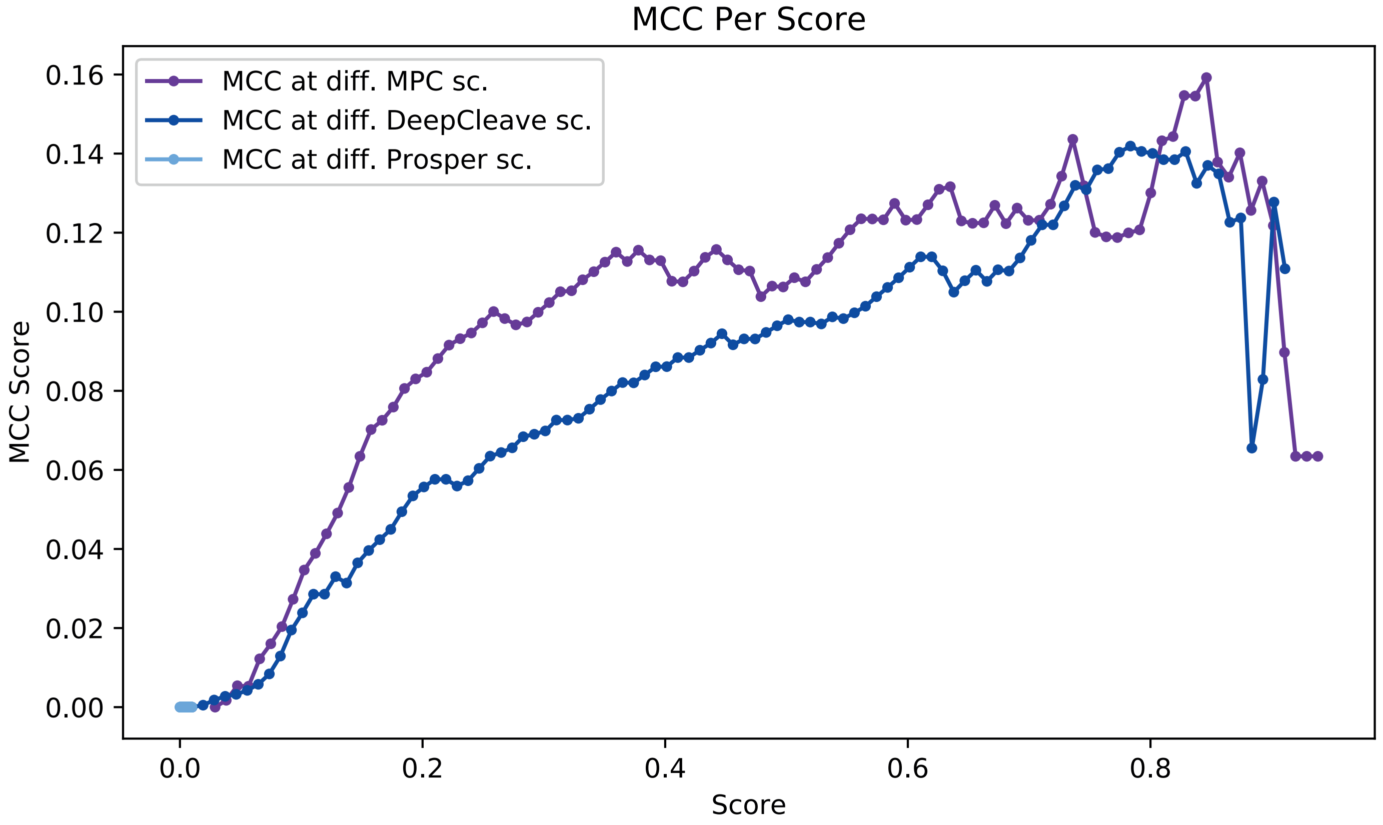


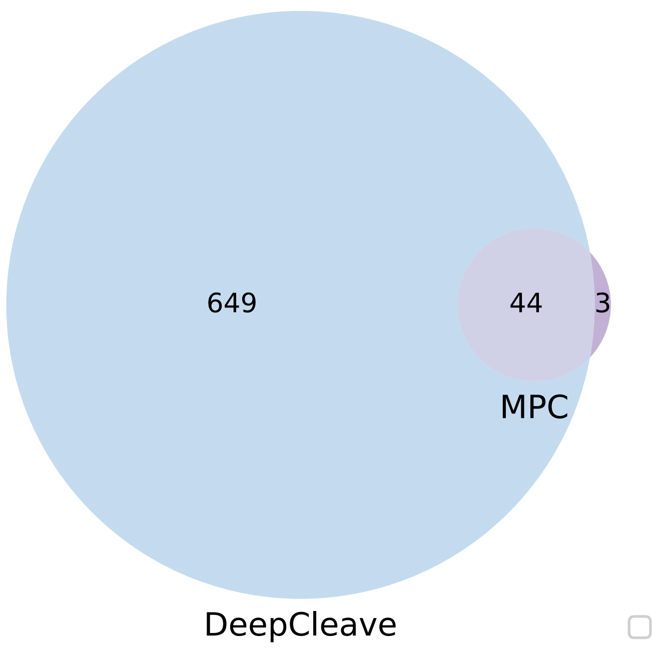

### MMP2 model performance against 15% test data

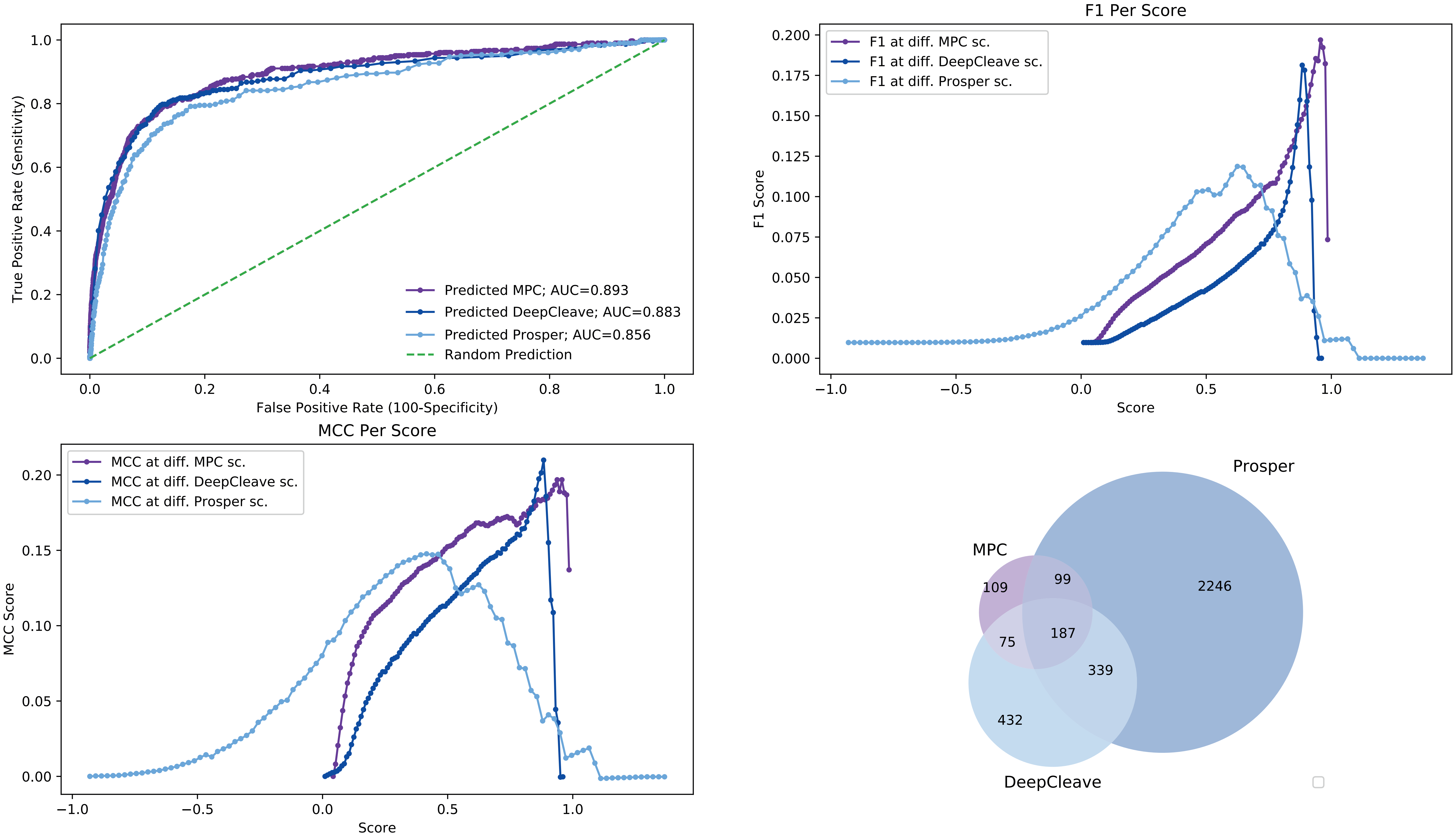

### MMP3 model performance against 15% test data

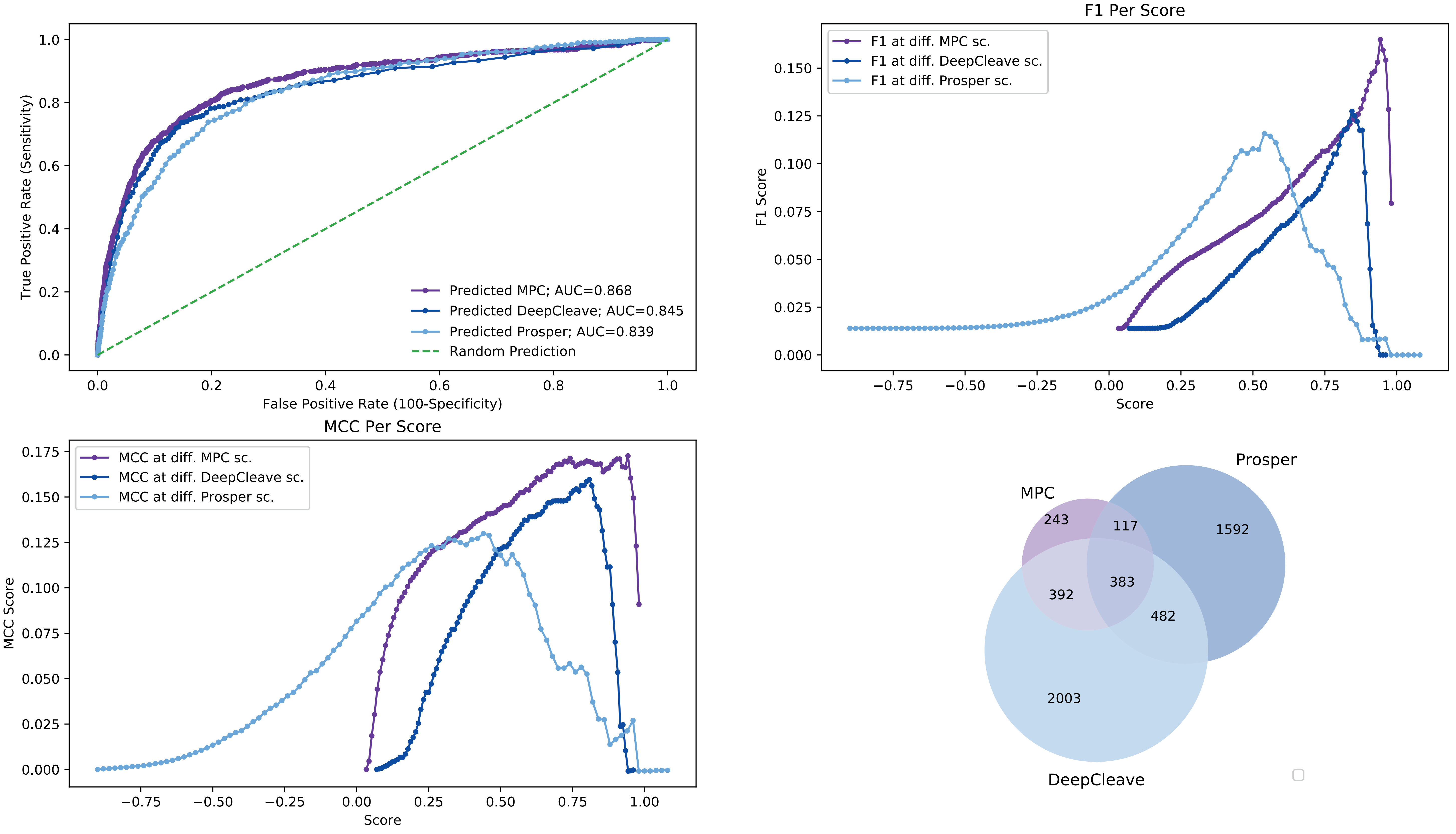

### MMP7 model performance against 15% test data

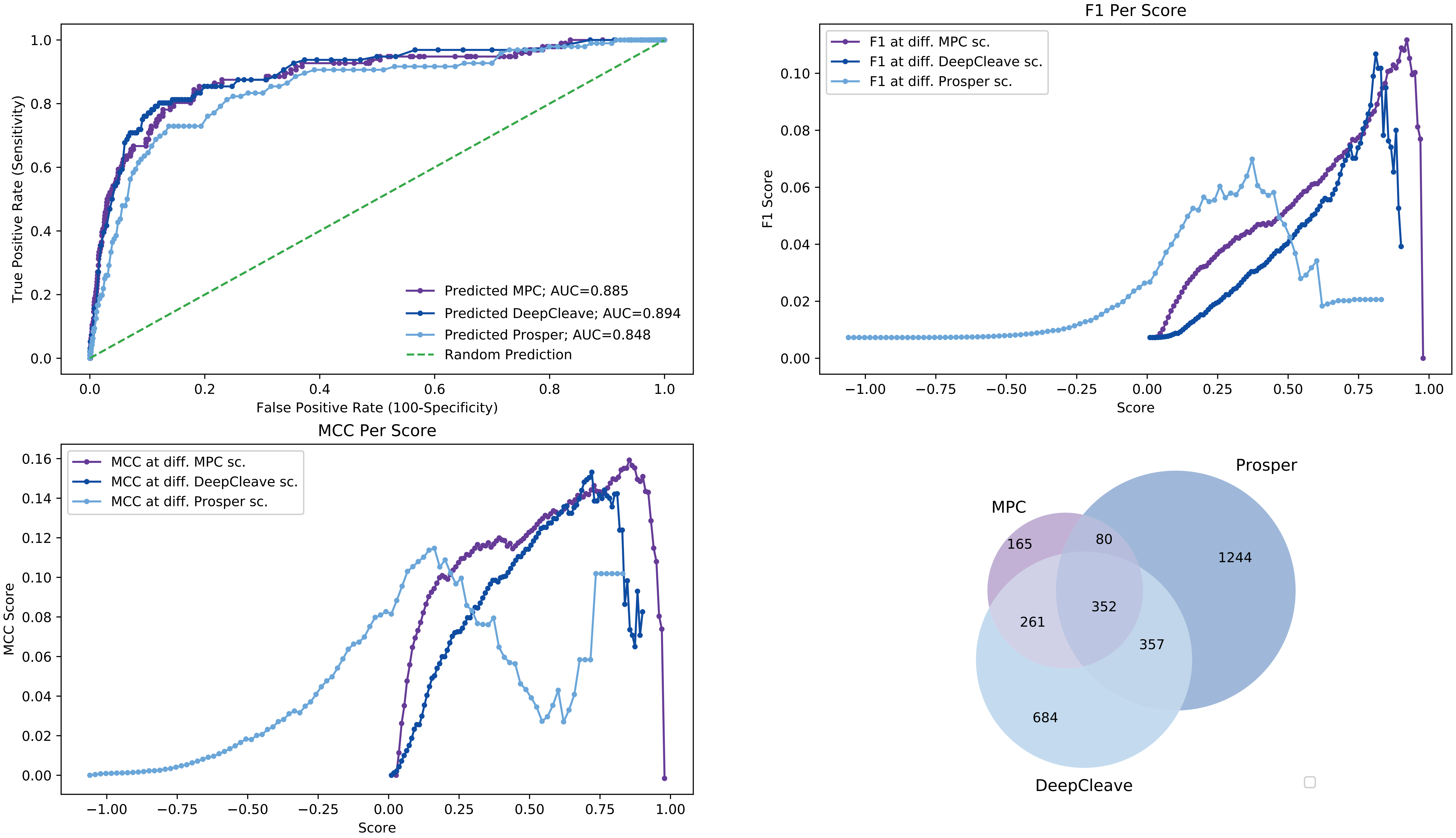

### MMP9 model performance against 15% test data

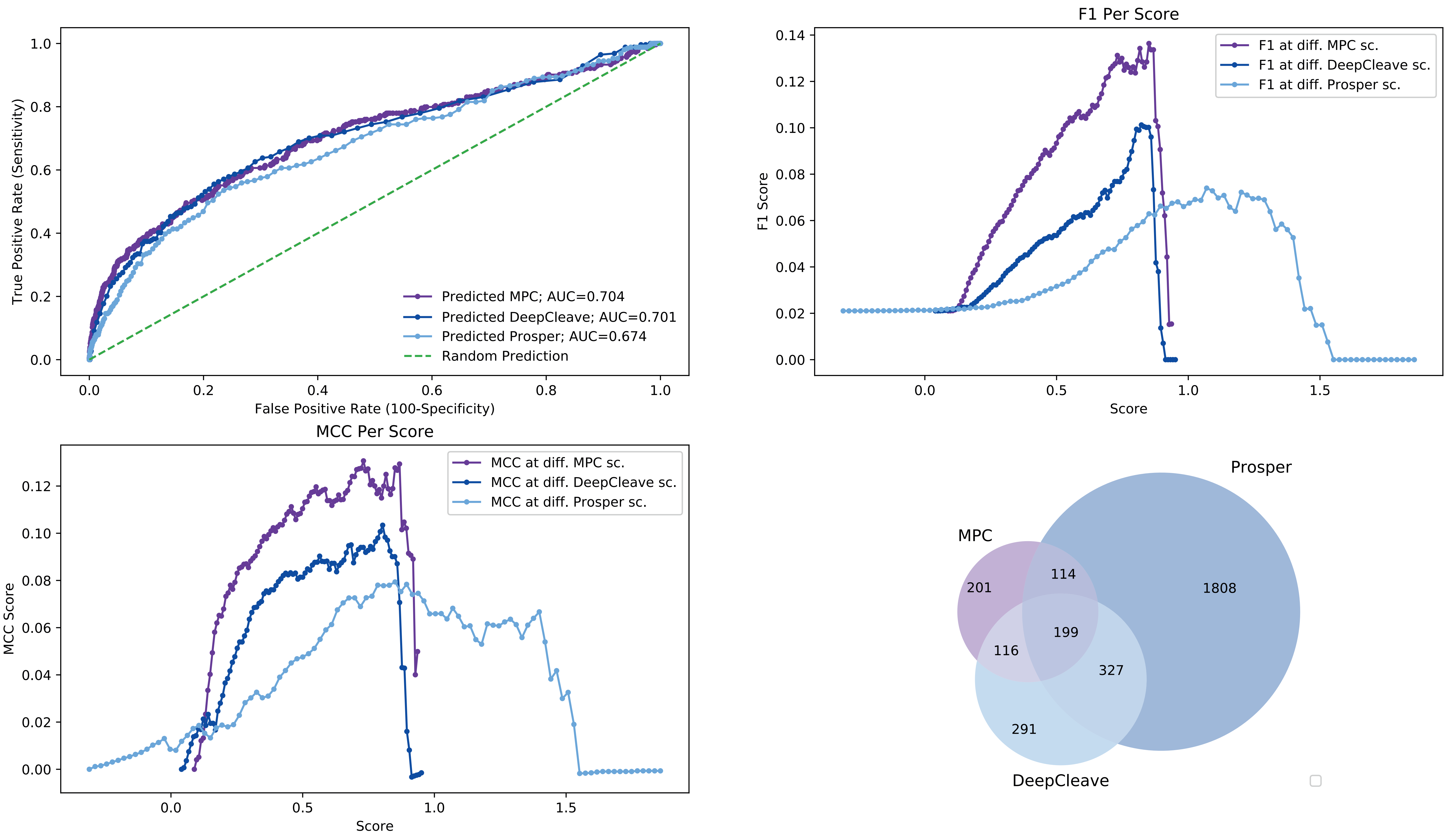

### MMP12 model performance against 15% test data

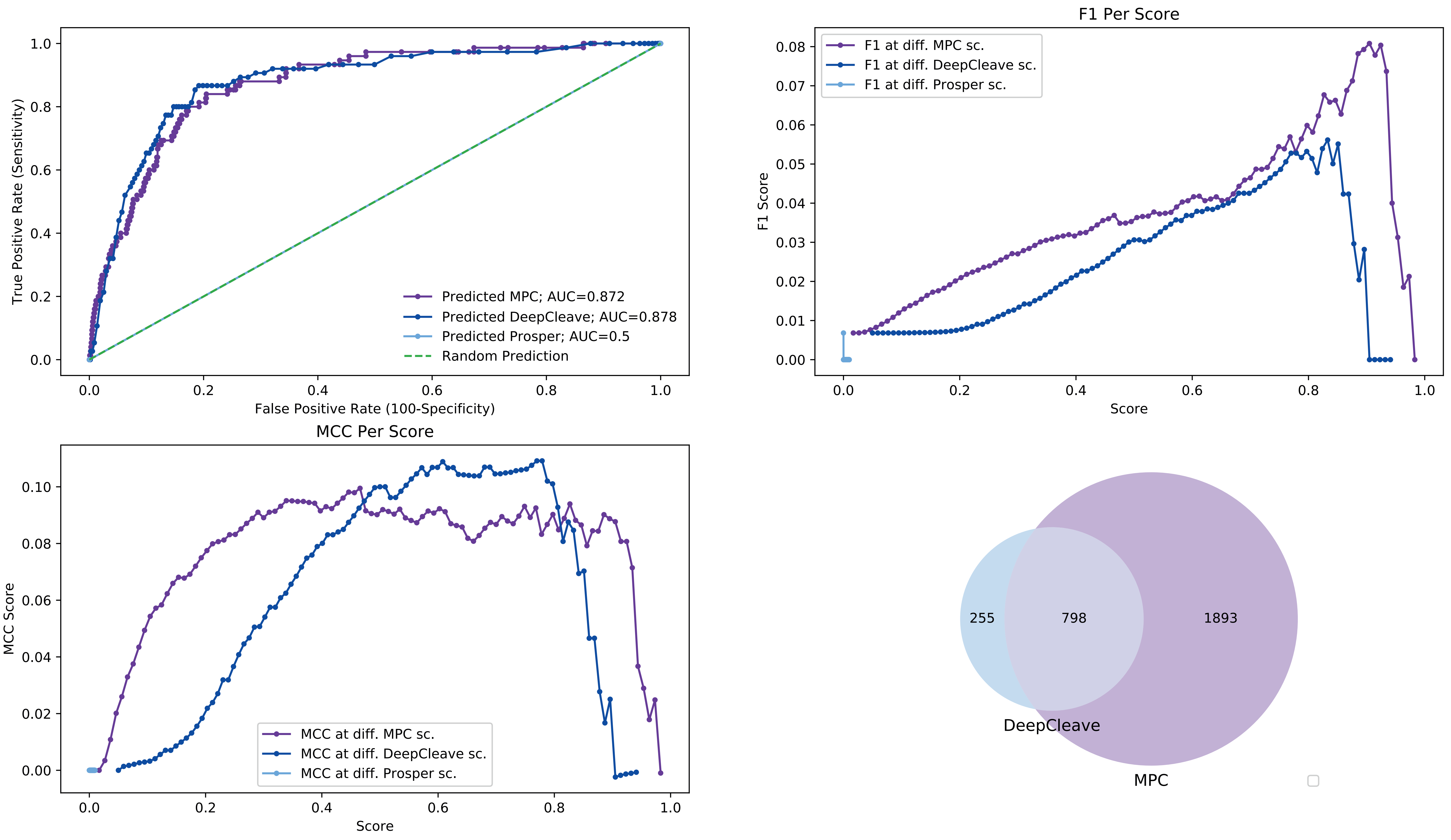

### MMP13 model performance against 15% test data

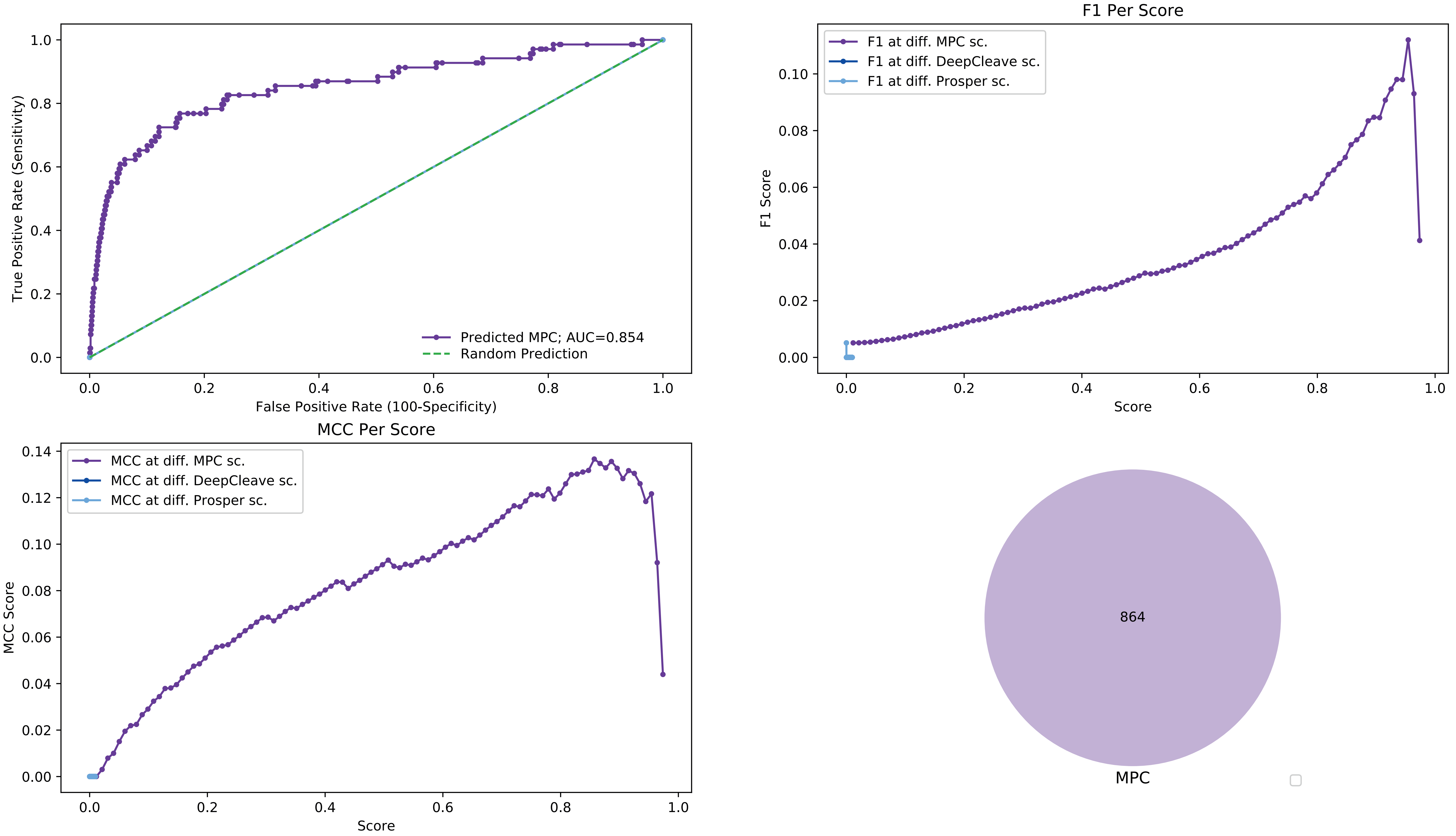

### MMP14 model performance against 15% test data

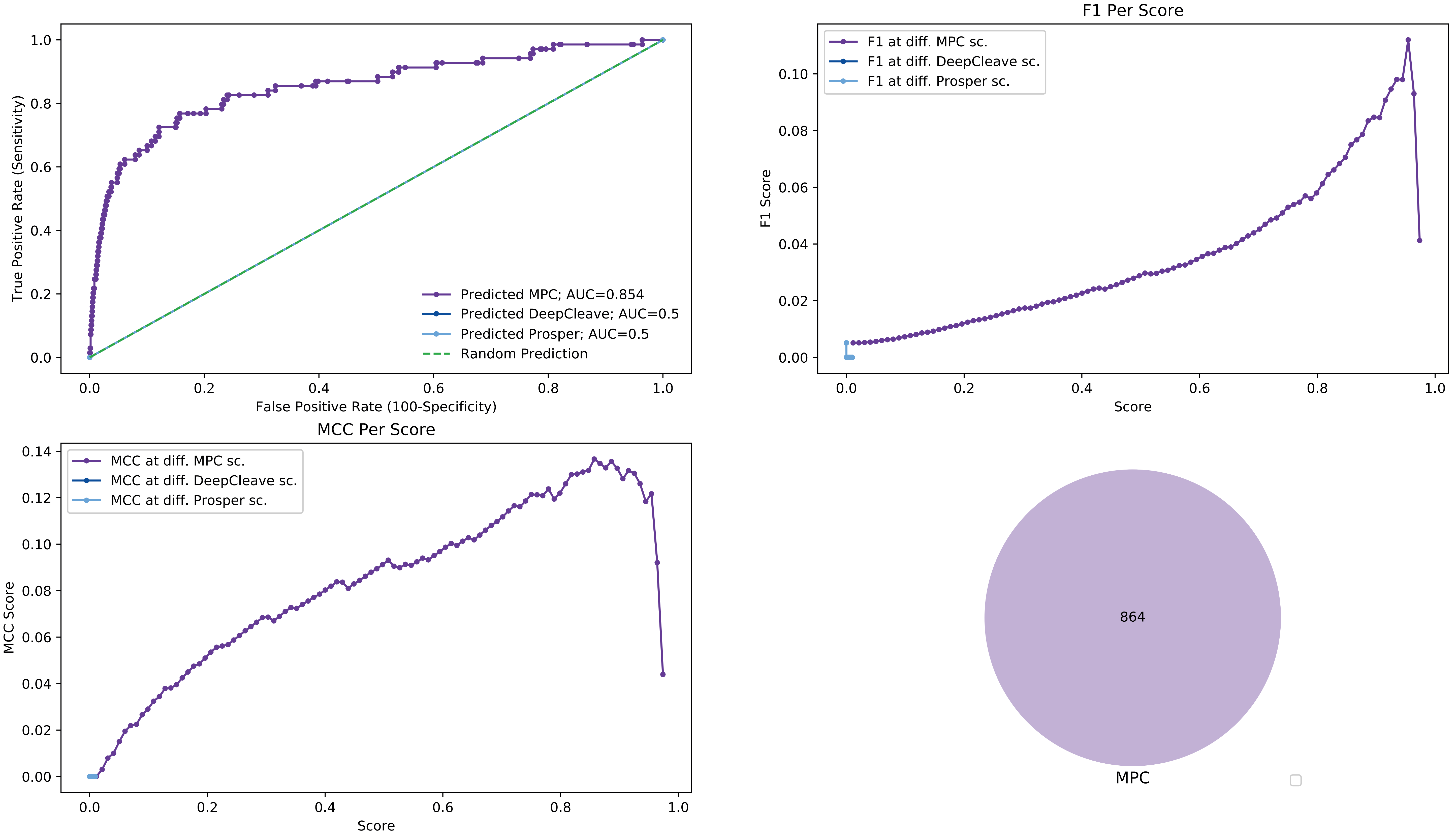

### Elastase-2 model performance against 15% test data

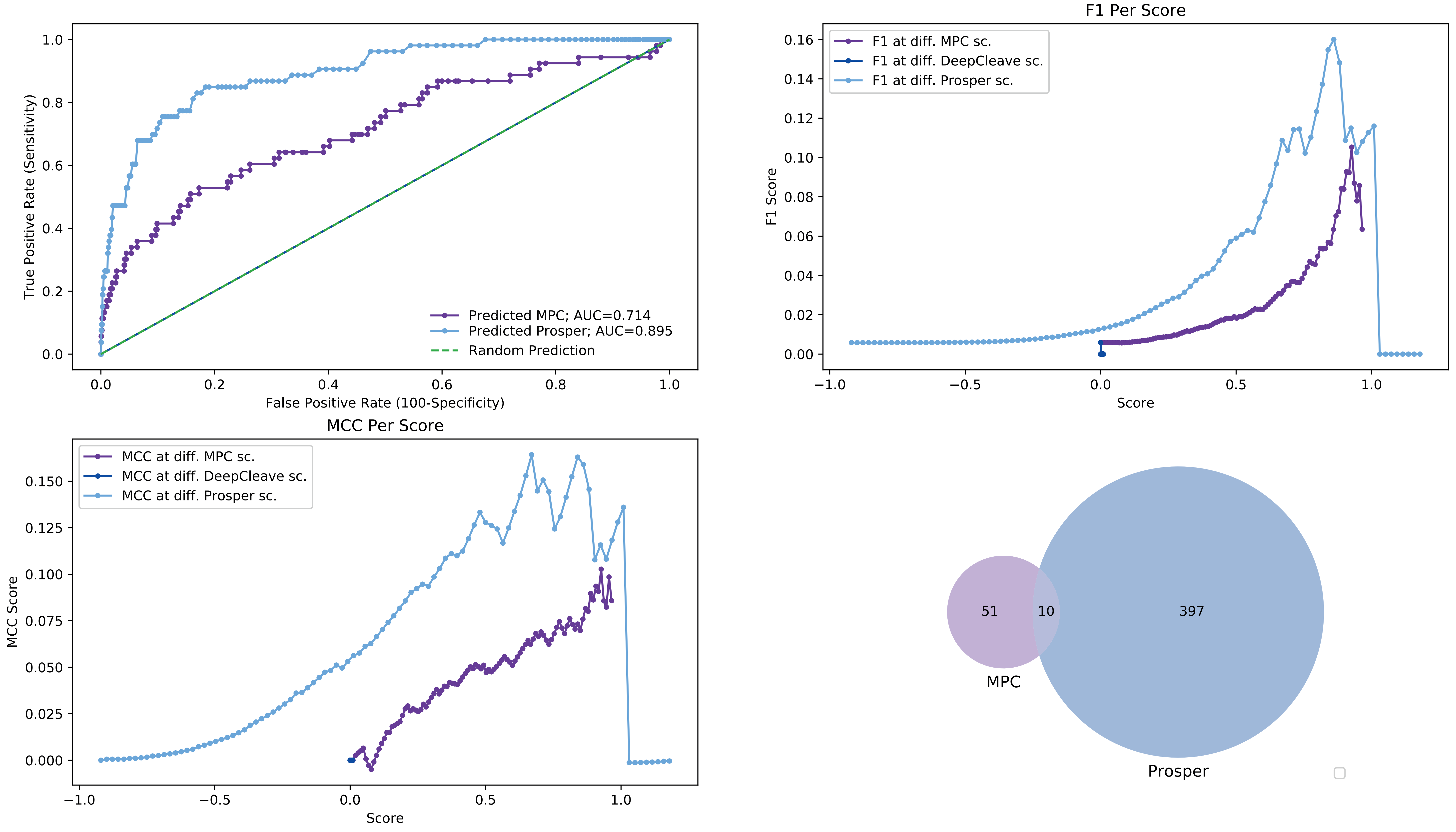

### Granzyme-B model performance against 15% test data

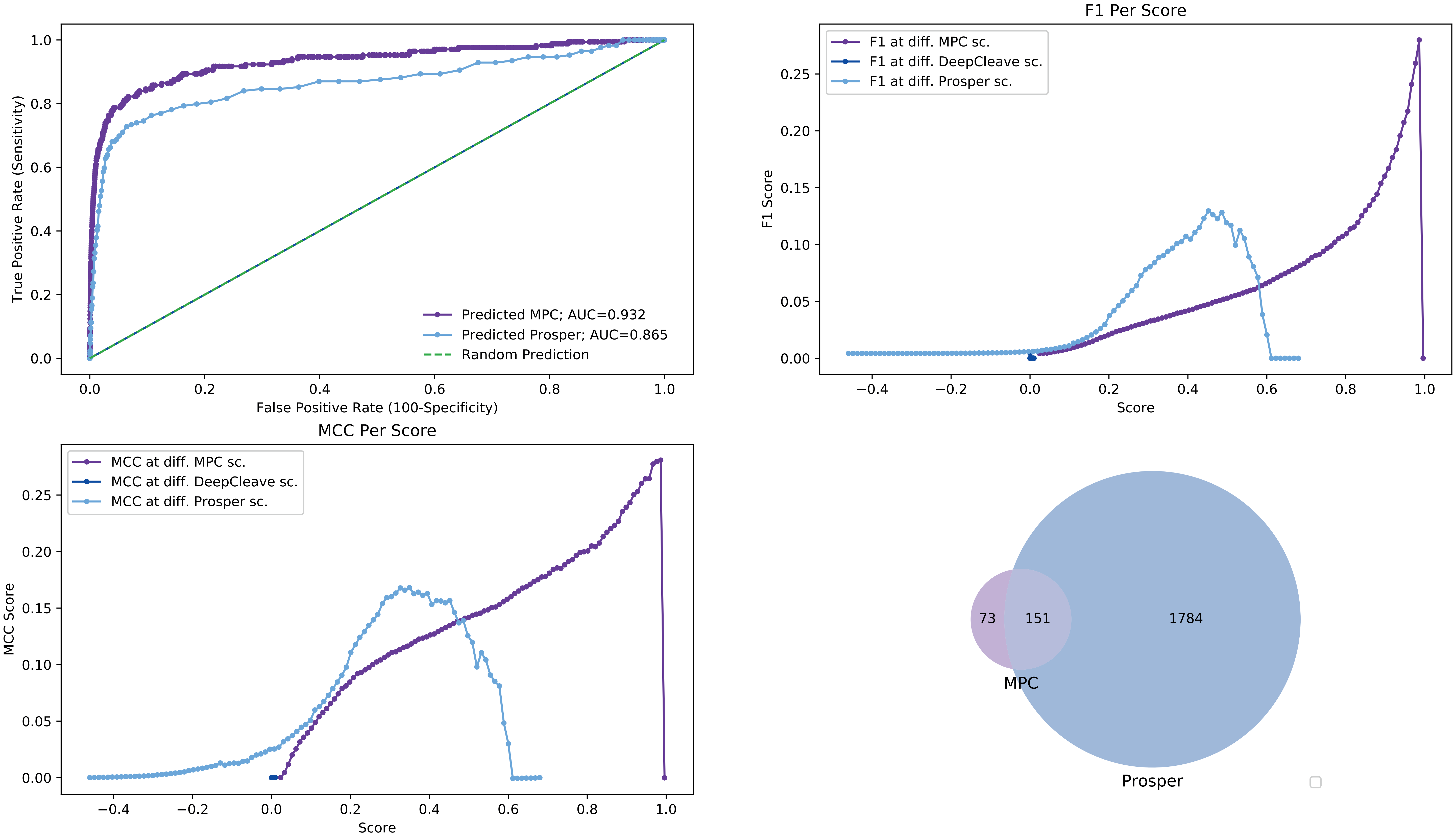

### Cathepsin-B model performance against 15% test data

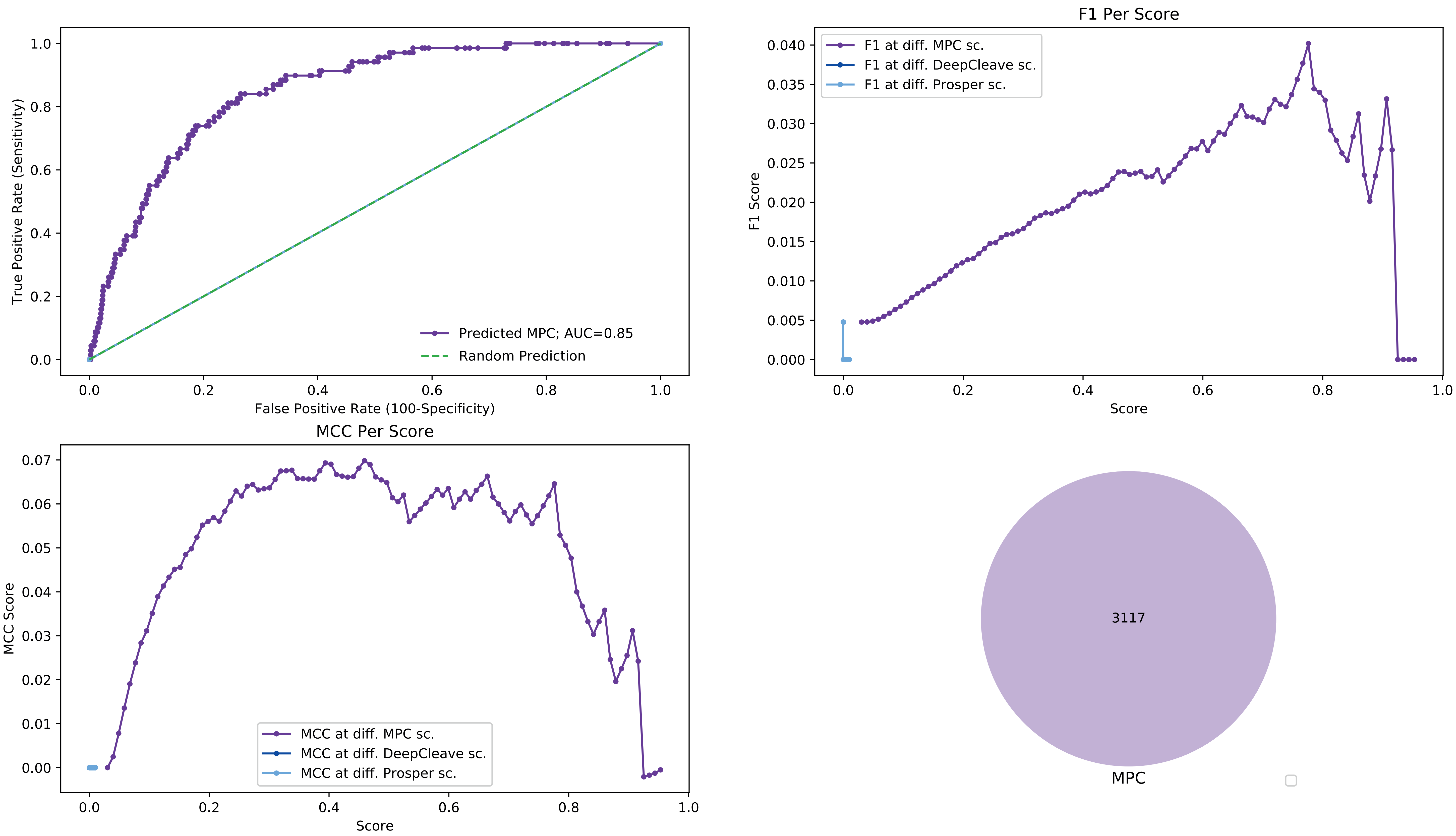

### Cathepsin-K model performance against 15% test data

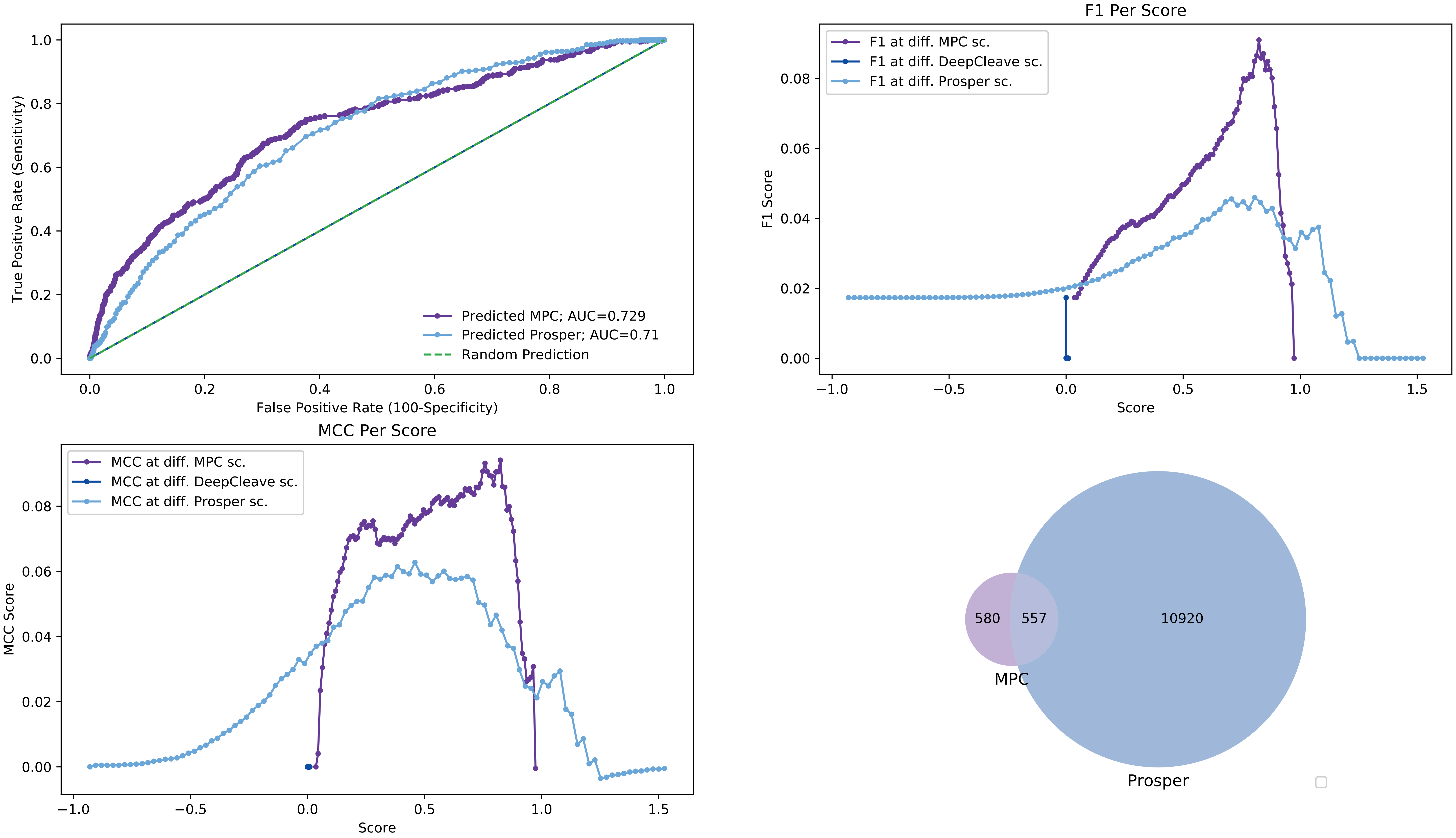

### Cathepsin-G model performance against 15% test data

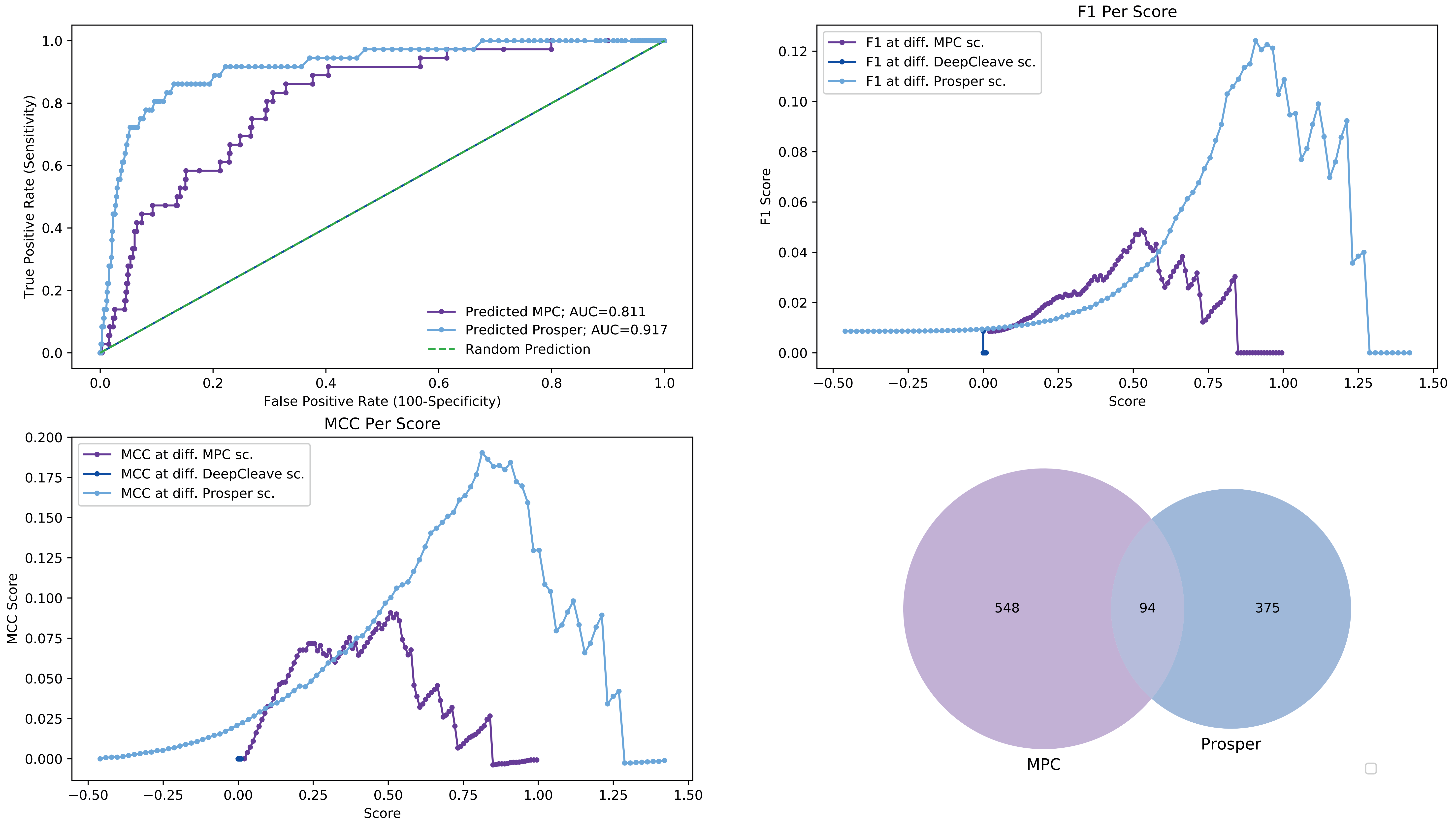

### Cathepsin-D model performance against 15% test data

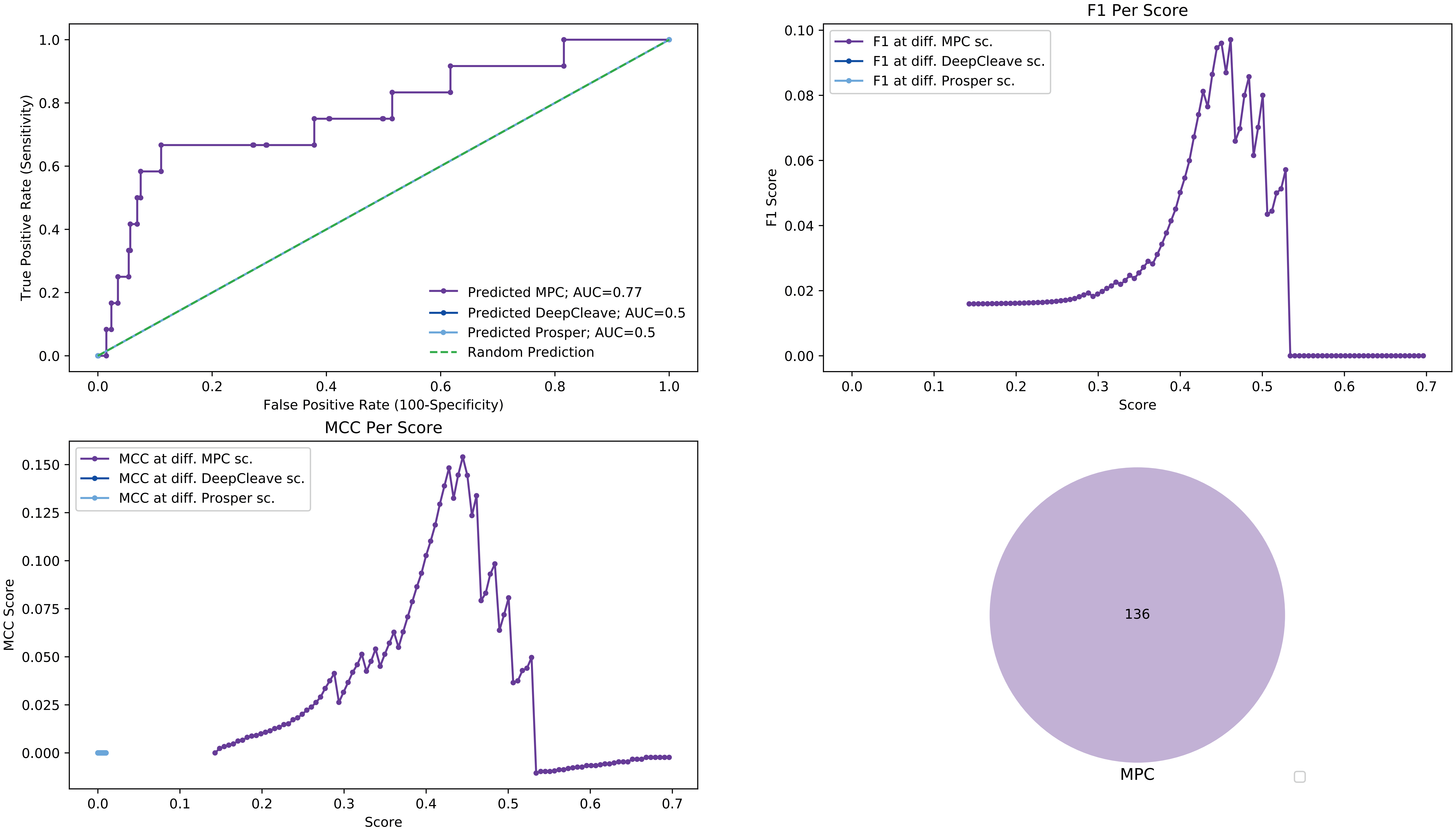
